# Laminin switches terminal differentiation fate of human trophoblast stem cells under chemically defined culture conditions

**DOI:** 10.1101/2021.09.30.462667

**Authors:** Victoria Karakis, Thomas McDonald, Abigail Cordiner, Adam Mischler, Adriana San Miguel, Balaji M Rao

## Abstract

Human trophoblast stem cells (hTSCs) have emerged as a powerful tool to model early placental development in vitro. Analogous to the epithelial cytotrophoblast in the placenta, hTSCs can differentiate into cells of the extravillous trophoblast (EVT) lineage or the multinucleate syncytiotrophoblast (STB). Here we present a chemically defined culture system for STB and EVT differentiation of hTSCs. Notably, in contrast to current approaches, we do not utilize transforming growth factor-beta inhibitors or a passage step for EVT differentiation, or forskolin for STB formation. Strikingly, under these conditions, presence of a single additional extracellular cue – lam-inin-1 – switched the terminal differentiation of hTSCs from STB to the EVT lineage. Activation of the sphingosine-1 receptor 3 receptor (S1PR3) using a chemical agonist could drive EVT differentiation of hTSCs in the absence of exogenous laminin, albeit less efficiently. To illustrate the utility of a chemically defined culture system for mechanistic studies, we examined the role of protein kinase C (PKC) signaling during hTSC differentiation to the EVT lineage. Inhibition of PKCα/β signaling significantly reduced HLA-G expression and the formation of HLA-G^+^ mesenchymal EVTs during hTSC differentiation mediated by laminin exposure; however, it did not prevent commitment to the EVT lineage or STB differentiation. The chemically defined culture system for hTSC differentiation established herein facilitates quantitative analysis of heterogeneity that arises during hTSC differentiation, and will enable mechanistic studies in vitro.

**Significance:** Despite its importance to a healthy pregnancy, early human placental development remains poorly understood. Mechanistic studies are impeded by restrictions on research with human embryos and fetal tissues, and significant differences in placentation between humans and commonly used animal models. In this context, human trophoblast stem cells (hTSCs) have emerged as attractive in vitro models for the epithelial cytotrophoblast of the early gestation human placenta. Here we describe chemically defined culture conditions for differentiation of hTSCs to the two major differentiated cell types – extravillous trophoblast and syncytiotrophoblast. These culture conditions enable in vitro studies to reveal molecular mechanisms regulating hTSC differentiation.

## Introduction

The placenta is a complex fetal organ with a vast network of villi that ensures efficient exchange of nutrients and waste across the maternal-fetal interface. Epithelial cytotrophoblasts (CTBs) of the early human placenta give rise to all trophoblast cell types in the placenta^1–4^. CTBs undergo cell fusion to form the multinucleate syncytiotrophoblast (STB) that overlays the CTB layer of placental villi^1,3^. The STB is subsequently bathed in maternal blood at ∼ 10 weeks of gestation, when blood flow to the placenta is established^5–7^. CTBs of placental villi anchored to the maternal decidua push through the syncytial layer and differentiate to extravillous trophoblasts (EVTs), first forming proliferative column trophoblasts adjacent to the villus tip^1,3^. At the distal end, column trophoblasts undergo an epithelial-to-mesenchymal transition (EMT) to become mature mesenchymal EVTs that invade the maternal decidua and parts of the myometrium^8–11^. These invasive trophoblasts aid in remodeling maternal spiral arteries and play a critical role in establishing sufficient perfusion of the placenta with maternal blood^12–15^.

Many pregnancy complications including preeclampsia, fetal growth restriction, miscarriage, and stillbirth, are often a result of impaired arterial remodeling and evidence points to improper EVT differentiation and invasion as a major contributor to these pathologies^3,16–19^. For instance, quantitative analyses of images from preeclamptic placentae biopsies demonstrate shallow trophoblast invasion compared with healthy placenta^20^. Previous studies have also suggested that increased apoptosis and an inability of CTBs to differentiate towards the EVT lineage may underlie shallow trophoblast invasion in preeclampsia^21,22^. Yet, despite the importance of EVT differentiation in placental health and pathology, molecular mechanisms underlying EVT differentiation and maturation to mesenchymal invasive EVTs remain poorly understood.

Mechanistic studies on early human placental development have been impeded primarily due to lack of suitable model systems. There are substantial restrictions on research with fetal tissue and human embryos and significant differences between placental development in common experimental animals and humans^23–28^. Additionally, there are significant differences in the transcriptome profiles of immortalized cell lines and primary trophoblasts^29^. Further, immortalized cell lines do not model the heterogeneity of cell types observed during EVT differentiation in vivo^30^. In this context, human trophoblast stem cells (hTSCs) derived from early gestation primary placental samples and blastocysts have emerged as powerful in vitro models of early human placental development^31^. hTSCs, which model the CTB in vivo, can be maintained in cell culture and can differentiate into STB or EVTs. In recent work, others and we have also shown that hTSCs can be derived from human pluripotent stem cells^32–40^, raising the exciting prospect of investigating pathological trophoblast development using induced pluripotent stem cells derived from somatic tissue (e.g. placenta) obtained at birth^41^. Nevertheless, current protocols for hTSC differentiation to EVT and STB limit mechanistic studies.

Most current protocols for in vitro differentiation include transforming growth factor-beta (TGFβ) inhibitors during EVT differentiation^31,38,39,42^. However, this is problematic because TGFβ plays an important role in EVT differentiation in vivo, and dysregulation of TGFβ signaling can result in placental pathologies^11,17,43–49^. Inhibition of TGFβ during EVT differentiation precludes investigation into the role of TGFβ signaling during normal or pathological trophoblast differentiation. Similarly, STB differentiation in vitro predominantly relies on the use of forskolin to induce cell fusion. Forskolin raises cyclic AMP concentrations causing upregulation of GCM1, which controls expression of fusion genes, syncytin-1 and −2^50^. The use of forskolin impedes studies on extracellular cues regulating STB differentiation. Finally, it is important to note that heterogeneity of cell types is a central feature of EVT differentiation in vivo. Initially, epithelial CTBs form proliferative proximal cell columns that express column markers MYC, VE-cadherin, Notch1, as well as CTB markers EGFR and ITGA6^3,8,11,31,45,51–60^. Differentiating EVTs at the distal end of the epithelial column remain proliferative; however, they begin to express ITGA5 and low levels of the mature EVT marker HLA-G, and lose CTB marker expression^52,57,59,61^. Subsequently, when these distal EVTs undergo EMT, they gain expression of ITGA1, express higher levels of HLA-G, and lose expression of markers that characterize the column trophoblasts^3,62^. Current protocols for EVT differentiation that include a passage step do not capture EVT heterogeneity and the sequential nature of CTB differentiation as mature mesenchymal EVTs are formed.

Here we present chemically defined culture conditions for differentiation of placenta- and hiPSC-derived hTSCs to EVTs and STB. Notably, our conditions do not involve a passage step, and exclude forskolin and TGFβ inhibition during STB and EVT differentiation, respectively. Using these culture conditions, we identified extracellular cues provided by laminin or activation of sphin-gosine-1 kinase receptor 3 (S1PR3) as critical inputs that switch differentiation hTSCs from STB to the EVT lineage.

## Results

### Chemically defined conditions for STB differentiation in the absence of forskolin

Placenta-derived CT29 and CT30 hTSCs and hiPSC-derived SC102A-1 hTSCs were cultured in trophoblast stem cell medium (TSCM) as described previously^31,37^. Differentiation was induced by passaging hTSCs into a defined trophoblast differentiation medium (DTDM) supplemented with epidermal growth factor (EGF) and the ROCK inhibitor, Y-27632, at passage for two days, and culturing them for an additional 4 days in DTDM (**Fig. 1A**). Upon passage, we initially observed an increase in cell number, but by day 6, cells in a flat monolayer without visible cell boundaries was observed (**Fig. S1A**); lacunae were also detected (**Fig. S1A**). Immunofluorescence revealed expression of STB markers, hCG, SDC-1, as well as the pan-trophoblast marker KRT7 on day 6 (**Figs. 1B, S1B, C**). Differentiated cells also expressed epidermal growth factor receptor (EGFR; **Figs. 1B, S1B, C**). EVT markers, HLA-G, Notch1, and ErbB2 as well as the CTB marker, p63, were not expressed, suggesting that EVT differentiation did not occur under these conditions (**Figs. 1B, S1B, C**). High-throughput qPCR analysis revealed that STB genes *CYP19A1, HSD3B1, ERVFRD1, ERVW1*, and *CSH1* were all upregulated by day 4 of STB differentiation in CT30 hTSCs and continued to be overexpressed on day 6 of differentiation compared to undifferentiated hTSCs at day 0 (**Fig. 1C**). The comprehensive dataset from qPCR analysis is included in **Tables S1 and S2**. Similarly, in CT29 hTSCs, *HSD3B1, ERVFRD1*, and *CSH1* were upregulated by day 4 of STB differentiation compared with hTSCs at day 0, in addition to the other STB markers *CGB, GCM1, and SDC1* (**Fig. S1D**). *CYP19A1* and *ERVW1* were upregulated on day 4 compared to day 2 of differentiation in CT29 hTSCs and *ERVW1* in CT30 hTSCs and *CGB* in CT29 hTSCs were even upregulated to a greater extent by day 6 of differentiation compared to day 4 (**Fig. 1C, Fig. S1D**); this shows an overall increase in STB gene expression over the 6-day differentiation period. On the other hand, CTB genes were downregulated over the 6-day differentiation period. Specifically, in CT30 hTSCs, *TP63, ELF5*, and *HAND1* were downregulated by day 2 of STB differentiation and *ITGA6* and *TEAD4* were downregulated by day 4 of STB differentiation compared to day 0 (**Fig. 1C**). *TP63, HAND1, ITGA6*, and *TEAD4* all had a significant reduction in expression on day 4 compared to day 2 of differentiation and on day 6 compared to day 4 of differentiation (**Fig. 1C**). In CT29 hTSCs, *TP63, ELF5, HAND1, ITGA6, TEAD4, CDH1* (E-Cadherin), and *YAP* were all downregulated by day 6 of STB differentiation with respect to day 0 (**Fig. S1D**). Lastly, a membrane stain revealed multinucleate cells with a fusion index comparable to that obtained during STB differentiation using the protocol described by Okae et al. (**Figs. 1D, E, S1E, F**)^31^. These results show that efficient STB differentiation of hTSCs occurs under our chemically defined conditions, in the absence of forskolin.

**Figure 1:**
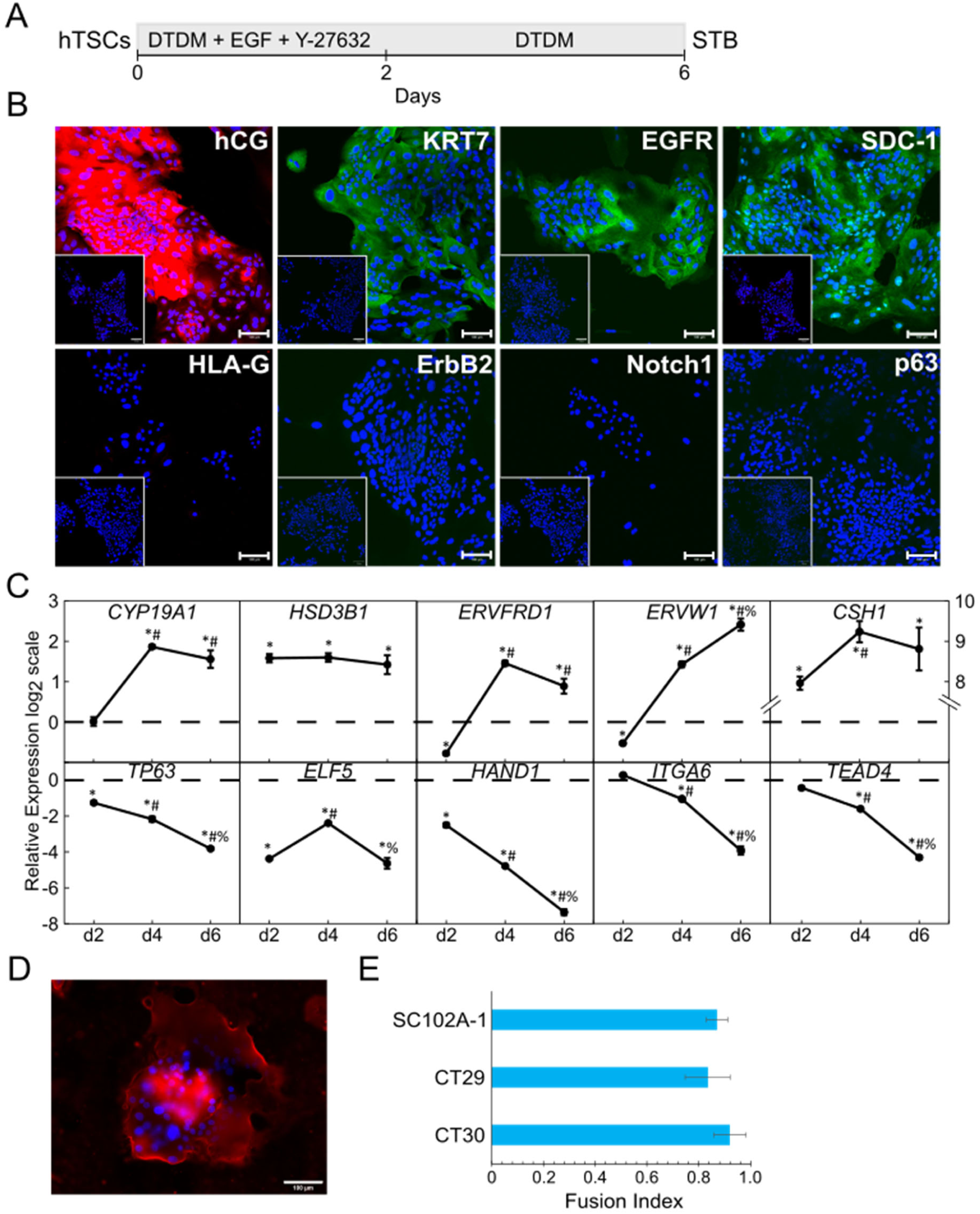
Chemically defined conditions for STB differentiation in the absence of for-skolin. (A) Schematic of protocol for hTSC differentiation to STB. (B) Confocal images of STB from CT30 hTSCs, staining for hCG, KRT7, EGFR, SDC-1, HLA-G, ErbB2, Notch1, p63. Nuclei were stained with DAPI. Inset is the respective isotype control. (C) Gene expression of *CYP19A1, HSD3B1, ERVFRD1, ERVW1, CSH1, TP63, ELF5, HAND1, ITGA6*, and *TEAD4* of STB from CT30 hTSCs. Three biological replicates were used. (Error bars, S.E., *p<0.05 for comparison with undifferentiated hTSCs (dashed line), ^#^p<0.05 for comparison with cells at day 2, ^%^p<0.05 for comparison with cells at day 4). (D) Fluorescent image of STB from CT30 hTSCs. Nuclei were stained with DAPI. Membrane was stained with Di-8-ANEPPS cell membrane stain. (E) Fusion efficiency of STB from CT30, CT29, and SC102A-1 hTSCs. Fusion index is calculated as (N-S)/T where N is the number of nuclei in the syncytia, S is the number of syncytia, and T is the total number of nuclei counted. (Error bars, S.D., n=3). Scale bars are 100µm for all images.

### Presence of laminin switches hTSC differentiation from STB to EVT fate

Okae et al. include Matrigel™ during EVT differentiation of hTSCs; absence of Matrigel™ results in formation of both EVT-like and STB-like cells^31^. Since laminin-1 is the major component of Matrigel™, we hypothesized that laminin-1 may mediate EVT differentiation of hTSCs^63^. Accordingly, we differentiated hTSCs using the previously described STB differentiation protocol (**Fig. 1A**), with the addition of laminin-1 for 2 days following passage, and further culture for an additional 4 days in DTDM (**Fig. 2A**). Addition of laminin resulted in a thin monolayer of matrix that solidified on the plate, covering the cells underneath (**Fig. S2A**). Upon initiation of differentiation, we observed that cells initially proliferated to form epithelial colonies. However, by day 6 of differentiation, a fraction of cells in these colonies underwent an epithelial-to-mesenchymal transition to form single, mesenchymal cells (**Figs. 2B, S2B**); this is reminiscent of mature mesenchymal EVTs forming from the distal end of trophoblast columns in vivo. Differentiated cells expressed the EVT markers HLA-G, Notch1, EGFR, VE-cadherin, CD9, ErbB2, and the pan-trophoblast marker KRT7; however, the CTB marker p63 was not expressed (**Figs. 2B, D, S2C, E, G, H**). In vivo, EVTs in the proximal column express Notch1 and lower levels of HLA-G whereas the later-stage mesenchymal EVT express HLA-G to a much greater extent and lose expression of Notch1^3,52,54,57,64,65^. Analogous to EVTs in vivo, cells in the epithelial colonies appeared to express higher levels of Notch1 and lower levels of HLA-G than the mesenchymal cells, which exhibited higher levels of HLA-G expression and lower Notch1 expression (**Figs. 2B, S2C, E**). Quantitative analysis revealed that cells in the bottom quartile of HLA-G expression (HLA-G^+^ i.e. 25% of total cells that showed the lowest HLA-G expression intensity) indeed expressed Notch1 with a greater intensity than cells that were of the top HLA-G expression quartile (HLA-G^++++^ i.e. 25% of total cells that showed the highest HLA-G expression intensity) in CT29 and CT30 EVTs; however, increase in Notch1 intensity for HLA-G^+^ cells was not statistically significant in SC102A-1 hTSCs (**Figs. 2C, S2D, F**). Consistent with our image analysis, flow cytometry analysis showed that HLA-G expression increased from day 2 to day 6 whereas Notch1 expression increased from day 2 to day 4 but then decreased on day 6 (**Figs. 2E, S2I, J**). This is consistent with previous studies that have reported Notch1 expression in early stage EVTs that constitute the proximal column trophoblast and higher HLA-G expression in late stage mesenchymal EVTs ^3,52,54,57,64,66^. Additional qPCR analysis showed increased expression of other EVT markers throughout EVT differentiation. Specifically, in CT30 hTSCs, *ITGA5, MMP2*, and *CDH5* expression increased by day 4 of differentiation and *CD9* expression increased by day 6 of differentiation compared with undifferentiated hTSCs at day 0 (**Fig 2F**). In CT29 hTSCs, *ITGA5, ITGA1, CD9*, and *CDH5* expression increased by day 6 of differentiation compared with day 0 (**Fig. S2K**). *MYC* expression is associated with EVT cell columns^3,54^ and was initially upregulated on day 2 in CT30 hTSCs but then decreased back to levels comparable to hTSCs on day 0 by day 6. In CT29 hTSCs, *MYC* expression decreased by day 6 of EVT differentiation compared with day 0 hTSCs. (**Fig. S2K**). Collectively, these data show an overall increase in EVT-associated gene expression over the 6-day differentiation period. Concomitantly, similar to STB differentiation, expression of CTB markers decreased during the 6-day differentiation. In CT30 hTSCs *TP63, ELF5, HAND1, ITGA6*, and *TEAD4* expression decreased by day 2 of differentiation (**Fig. 2F**). In CT29 hTSCs, *TP63, ELF5, HAND1, CDH1*, and *TEAD4* were all downregulated by day 6 of EVT differentiation (**Fig. S2K**). The comprehensive dataset from qPCR analysis is included in **Tables S1 and S2**. Taken together, our results are consistent with a 2-stage EVT differentiation process where hTSCs initially commit to the EVT lineage and gain expression of column EVT markers (day 0 to 4). Subsequently, cells undergo EMT to form single-cell, mesenchymal EVTs that express high levels of HLA-G and other EVT markers, and decreased expression of Notch1 (day 4 to 6). Importantly, because the only difference between the STB and EVT differentiation protocol is the addition of laminin-1 during day 0 to day 2, we can conclude that laminin-1 switches the terminal trophoblast differentiation fate from STB to EVT.

**Figure 2:**
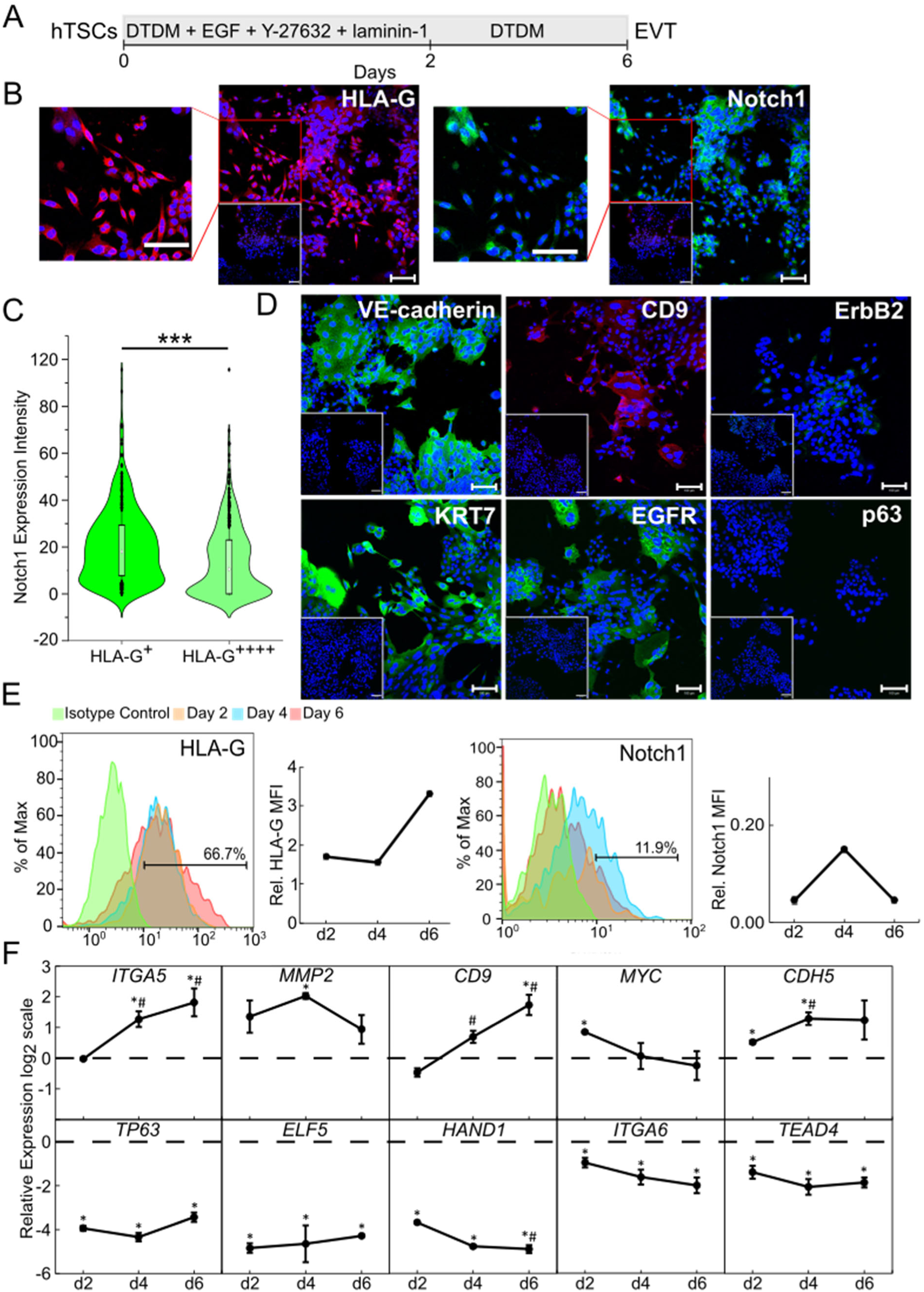
Presence of laminin switches hTSC differentiation from STB to EVT fate. (A) Schematic of protocol for hTSC differentiation to EVT. (B) Confocal images of EVT from CT30 hTSCs, staining for HLA-G and Notch1. Nuclei were stained with DAPI. Inset is the respective isotype control. Outcrop is the magnified image. (C) Quantitative analysis of Notch1 expression intensity of CT30 EVTs from the bottom (HLA-G^+^) and top (HLA-G^++++^) 25% of HLA-G expression intensity EVTs (n=550, each). Analysis was performed in MATLAB and at least 2 biological replicates were used. (***p<0.0005). (D) Confocal images of EVT from CT30 hTSCs, staining for VE-cadherin, CD9, ErbB2, KRT7, EGFR, and p63. Nuclei were stained with DAPI. Inset is the respective isotype control. (E) Flow cytometry histogram of HLA-G and Notch1 expression of CT30 EVTs on day 2, day 4, and day 6 compared to an isotype control and their relative mean fluorescence intensity (MFI). Relative MFI is calculated as (X-C)/C where X is the experimental MFI and C is the isotype control MFI. (F) Gene expression of *ITGA5, MMP2, CD9, MYC, CDH5, TP63, ELF5, HAND1, ITGA6*, and *TEAD4* of EVT from CT30 hTSCs. Three biological replicates were used. (Error bars, S.E., *p<0.05 for comparison with undifferentiated hTSCs (dashed line), ^#^p<0.05 for comparison with cells at day 2, ^%^p<0.05 for comparison with cells at day 4). Scale bars are 100µm for all images.

### Activation of S1PR3 agonist can mediate EVT differentiation of hTSCs in the absence of exogenous laminin

When laminin-1 was replaced by a small molecule agonist of S1PR3 (CYM5541) (**Fig. 3A**), we observed formation of Notch1^+^ colonies and single, HLA-G^+^ mesenchymal cells, suggesting that EVT differentiation occurred under these conditions (**Figs. 3B, S3A, B**). Differentiated cells obtained in the presence of the S1PR3 agonist showed upregulation of EVT markers *ITGA5, MMP2*, and *ITGA1* by day 4 of differentiation compared with undifferentiated hTSCs at day 0 (**Fig. 3C, S3C**). Additionally, in CT29 hTSCs, *MMP9* and *CDH5* were upregulated on day 4 and *CD9* was upregulated on day 6 of differentiation compared with day 0 (**Fig. S3C**). Expression of CTB markers *TP63, ELF5* and *HAND1* in CT30 hTSCs were downregulated by day 4 of differentiation and *ITGA6* and *TEAD4* were downregulated by day 6 compared to day 0 (**Fig. 3C**). Similarly, in CT29 hTSCs, *ELF5* and *HAND1* were downregulated by day 4 and *TEAD4* was downregulated by day 6 of differentiation (**Fig. S3C**). Notably, however, the expression intensity of HLA-G was lower in EVTs obtained by treatment with the S1PR3 agonist relative to EVTs obtained by differentiation in the presence of laminin across three cell lines (**Figs. 3D-F**). Collectively, these data showed that EVT differentiation of hTSCs could be triggered by activation of S1PR3 in the absence of laminin, albeit with lower efficiency.

**Figure 3:**
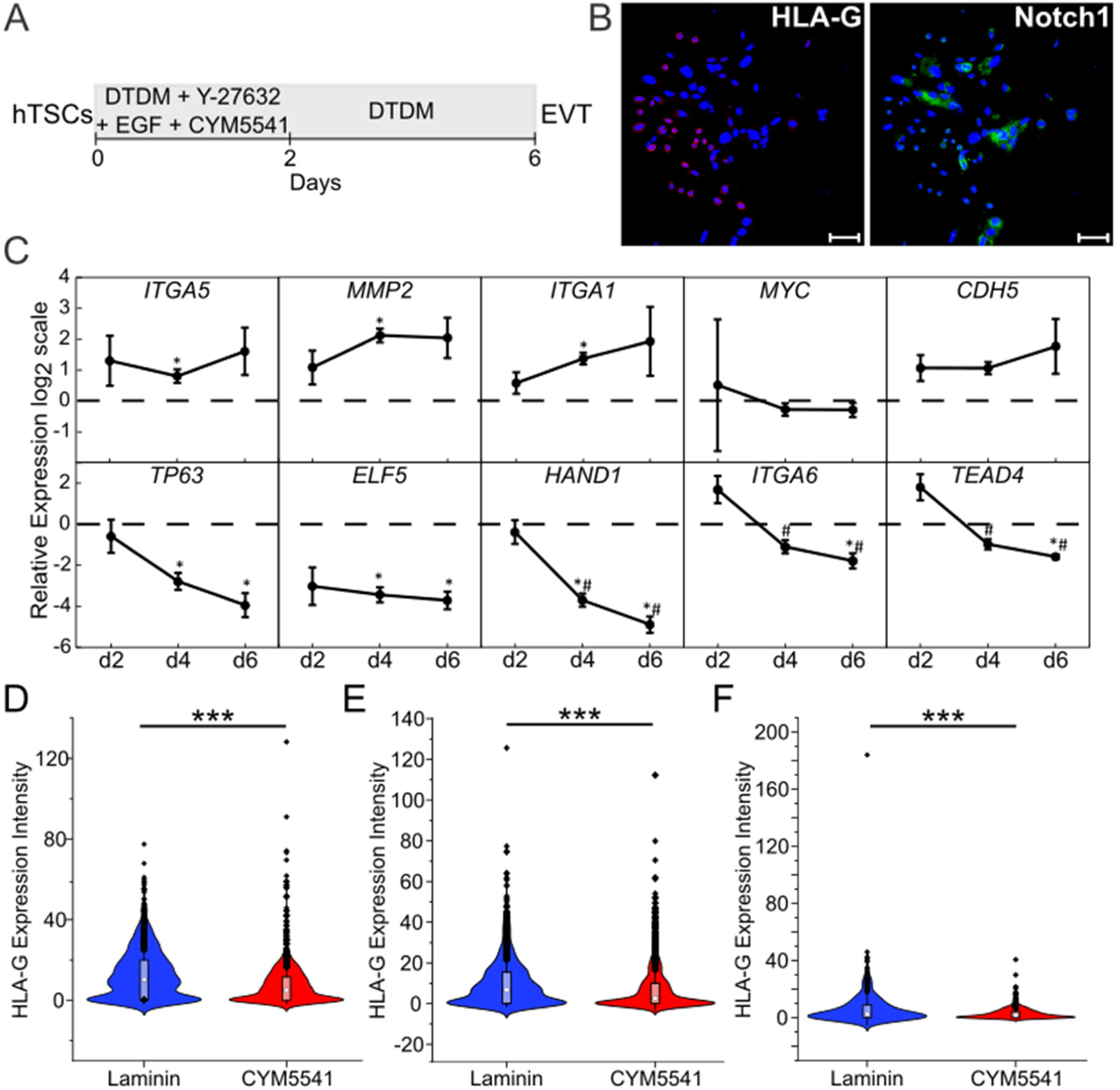
Activation of S1PR3 agonist can mediate EVT differentiation of hTSCs in the absence of exogenous laminin. (A) Schematic of protocol for hTSC differentiation to EVT obtained by exposure to a S1PR3 agonist, CYM5541. (B) Confocal images of EVT obtained by exposure to a S1PR3 agonist, CYM5541 from CT30 hTSCs, staining for HLA-G and Notch1. Nuclei were stained with DAPI. (C) Gene expression of *ITGA5, MMP2, ITGA1, MYC, CDH5, TP63, ELF5, HAND1, ITGA6*, and *TEAD4* of EVT obtained by exposure to a S1PR3 agonist, CYM5541 from CT30 hTSCs. Three biological replicates were used. (Error bars, S.E., *p<0.05 for comparison with undifferentiated hTSCs (dashed line), ^#^p<0.05 for comparison with cells at day 2, ^%^p<0.05 for comparison with cells at day 4). (D) Quantitative analysis of HLA-G expression intensity in CT30 EVTs obtained by exposure to laminin (n=2197) or S1PR3 agonist, CYM5541 (n=2736). Analysis was performed in MATLAB and at least 2 biological replicates were used. (***p<0.0005). (E) Quantitative analysis of HLA-G expression intensity in CT29 EVTs obtained by exposure to laminin (n=2592) or S1PR3 agonist, CYM5541 (n=4398). Analysis was performed in MATLAB and at least 2 biological replicates were used. (***p<0.0005). (F) Quantitative analysis of HLA-G expression intensity in SC102A-1 EVTs obtained by exposure to laminin (n=498) or S1PR3 agonist, CYM5541 (n=749). Analysis was performed in MATLAB and at least 2 biological replicates were used. (***p<0.0005). Scale bars are 100µm for all images.

### Inhibition of PKCα/β adversely affects laminin-mediated EVT differentiation

Our studies showed that exposure to laminin or activation of S1PR3 could trigger EVT differentiation of hTSCs. Protein kinase C (PKC) α/β has been reported to be downstream of integrin- and S1PR-mediated signaling^67–71^. Therefore, we investigated the effect of PKCα/β inhibition on EVT differentiation, using the small molecule inhibitor Gö6976. Specifically, we inhibited PKCα/β during laminin exposure (**Fig. 4A**). Strikingly, we observed that single, mesenchymal HLA-G^+^ EVTs did not form in the presence of PKCα/β inhibition (**Figs. 4B, S4A, B**). We also observed significant cell death for hiPSC-derived hTSCs under these conditions. Few cells remained in culture even after 2x initial seeding densities were used; therefore, these cells were excluded from further analysis. Quantitative analysis of a comprehensive set of images showed that Notch1 intensity at day 6 was higher in the presence of PKCα/β inhibition (**Figs. 4C, S4C**). On the other hand, HLA-G intensity was significantly reduced (**Figs. 4D, S4D**). We further conducted qPCR analysis to examine markers other markers of EVT differentiation. We observed that expression of EVT markers *ITGA5* in CT30 hTSCs, *MYC* in CT29 hTSCs, and *MMP2, ITGA1*, and *CDH5* in both CT29 and CT30 hTSCs all were significantly lower on day 4 in the presence of PKCα/β inhibition relative to controls (**Fig. 4E, Fig. S4E**). Nevertheless, expression of *ITGA5, ITGA1*, and *CDH5* was upregulated by day 6 relative to day 0 hTSCs (**Fig. 4E, S4E**). Similarly, in CT30 hTSCs, the expression of CTB markers, *TP63, ELF5, HAND1* and *TEAD4* was higher at day 4 or day 6 of differentiation in the presence of PKCα/β inhibition relative to the laminin control (**Fig. 4E**). Nevertheless, CTB markers were downregulated relative to undifferentiated hTSCs at day 0 (**Figs. 4E, S4E**). Thus, PKCα/β inhibition did not prevent differentiation to the EVT lineage, although formation of mesenchymal HLA-G^+^ cells was greatly reduced. Furthermore, PKCα/β inhibition did not affect STB differentiation of hTSCs; expression of hCG and SDC-1 was unaffected by the presence of PKCα/β inhibition during STB differentiation (**Figs. 4F, G, S4F, G**). Finally, it is important to note that the PKCα/β inhibitor Gö6976 can also inhibit protein kinase D1 (PKD1)^72^. Therefore, to confirm that our results were not a consequence of PKD1 inhibition, we conducted hTSC differentiation to EVT and STB in the presence of a specific PKD1 inhibitor, CRT0066101 (**Figs. S5A, C**). We observed that EVT and STB differentiation remained unaffected in the presence of CRT0066101; specifically, we observed HLA-G^+^ single cells and Notch1^+^ epithelial colonies under EVT differentiation conditions, and expression of hCG and SDC-1 under STB differentiation conditions (**Figs. S5B, D**). Taken together, our results show that PKCα/β signaling is necessary for formation of HLA-G^+^ mesenchymal EVTs during differentiation mediated by laminin exposure; however inhibition of PKCα/β signaling does not prevent commitment to the EVT lineage nor STB differentiation.

**Figure 4:**
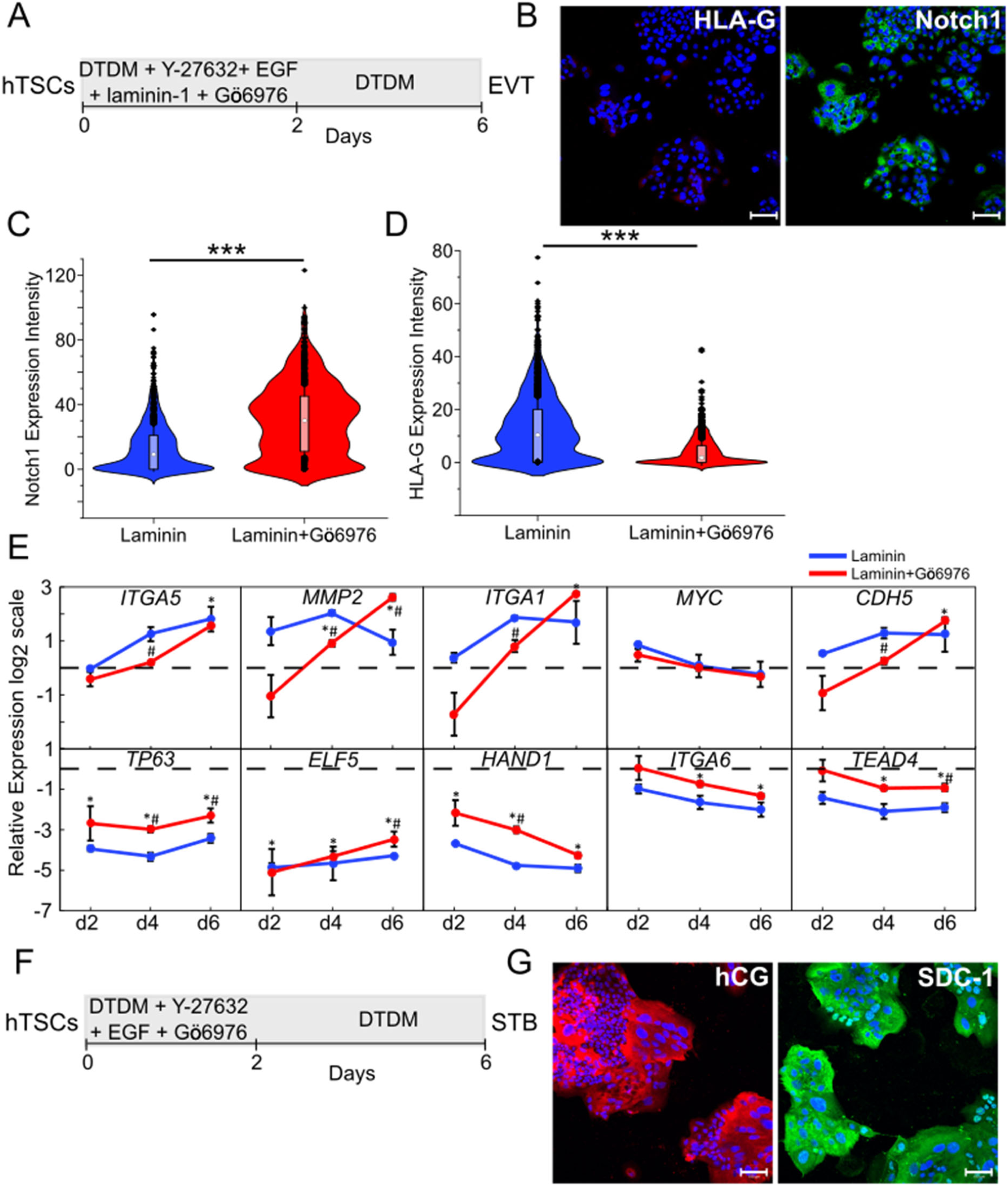
Inhibition of PKCα/β adversely affects laminin-mediated EVT differentiation. (A) Schematic of protocol for hTSC differentiation to EVT obtained by exposure to laminin in the presence of a PKCα/β inhibitor, Gö6976. (B) Confocal image of EVT obtained by exposure to laminin in the presence of a PKCα/β inhibitor, Gö6976 from CT30 hTSCs, staining for HLA-G and Notch1. Nuclei were stained with DAPI. (C) Quantitative analysis of Notch1 expression intensity in CT30 EVTs obtained by exposure to laminin (n=2197) or laminin in the presence of a PKCα/β inhibitor, Gö6976 (n=2204). Analysis was performed in MATLAB and at least 2 biological replicates were used. (***p<0.0005). (D) Quantitative analysis of HLA-G expression intensity in total CT30 EVTs obtained by exposure to laminin (n=2197) or laminin in the presence of a PKCα/β inhibitor, Gö6976 (n=2204). Analysis was performed in MATLAB and at least 2 biological replicates were used. (***p<0.0005). Data for laminin is same as used in Figure 3. (E) Gene expression of *ITGA5, MMP2, ITGA1, MYC, CDH5, TP63, ELF5, HAND1, ITGA6*, and *TEAD4* of EVTs obtained by exposure to laminin in the absence (blue) and presence (red) of the PKCα/β inhibitor, Gö6976 from CT30 hTSCs. Three biological replicates were used. (Error bars, S.E., *p<0.05 for comparison with undifferentiated hTSCs (dashed line), ^#^p<0.05 for comparison with EVTs obtained by exposure with laminin at the same day). (F) Schematic of protocol for hTSC differentiation to STB in the presence of a PKCα/β inhibitor, Gö6976. (G) Confocal image of STB in the presence of a PKCα/β inhibitor, Gö6976 from CT30 hTSCs, staining for hCG and SDC-1. Nuclei were stained with DAPI. Scale bars are 100µm for all images.

### PKCα/β inhibition differentially affects EVT differentiation mediated by laminin exposure and S1PR3 activation

S1PR3 activation can mediate EVT differentiation in the absence of laminin, although less efficiently. We investigated the effect of PKCα/β inhibition on EVT differentiation mediated by S1PR3 activation (**Fig. 5A**). We observed formation of HLA-G^+^ mesenchymal EVTs and Notch1^+^ colonies under these conditions (**Fig 5B, S6A, B**). Quantitative image analysis revealed an increase in both HLA-G and Notch1 expression intensity at day 6 when PKCα/β was inhibited during S1PR3 agonist treatment (**Fig. 5C, D, S6C-F**). In contrast, PKCα/β inhibition during laminin exposure led to significant reduction in both HLA-G expression intensity and the formation of HLA-G^+^ mesenchymal cells. Thus, PKCα/β has differential effects on EVT differentiation mediated by laminin and S1PR3 agonist. However, as noted earlier, EVT differentiation mediated by S1PR3 agonist in the absence of exogenous laminin is less efficient; lower overall HLA-G expression is observed than in the case of laminin-mediated EVT differentiation.

**Figure 5:**
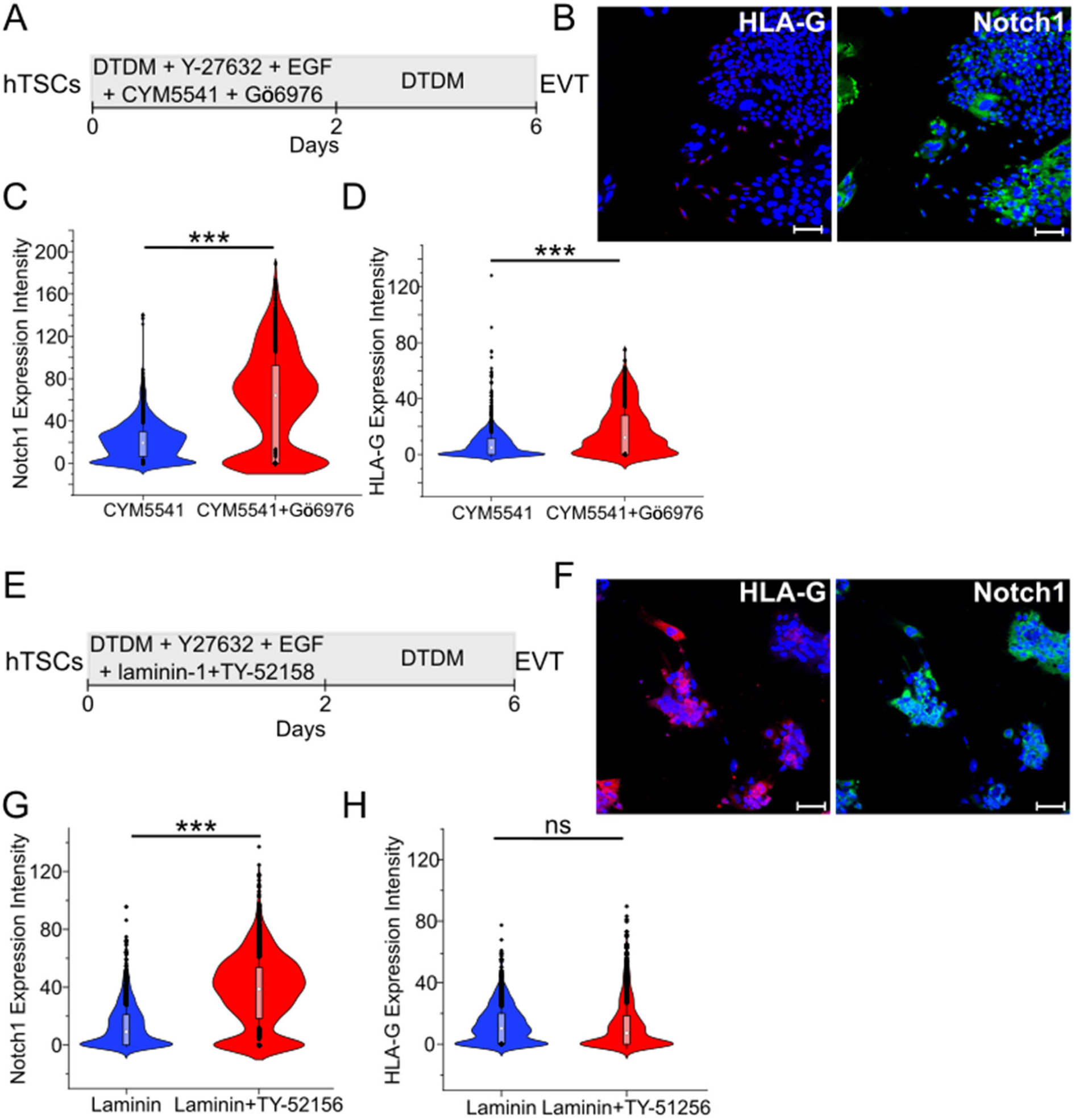
PKCα/β inhibition differentially affects EVT differentiation mediated by laminin exposure and S1PR3 activation. (A) Schematic of protocol for hTSC differentiation to EVT obtained by exposure to a S1PR3 agonist, CYM5541 in the presence of a PKCα/β inhibitor, Gö6976. (B) Confocal image of EVT obtained by exposure to a S1PR3 agonist, CYM5541 in the presence of a PKCα/β inhibitor, Gö6976 from CT30 hTSCs, staining for HLA-G and Notch1. Nuclei were stained with DAPI. (C) Quantitative analysis of Notch1 expression intensity in CT30 EVTs obtained by exposure to a S1PR3 agonist, CYM5541 (n=2736) or CYM5541 in the presence of a PKCα/β inhibitor, Gö6976 (n=1633). Analysis was performed in MATLAB and at least 2 biological replicates were used. (***p<0.0005). (D) Quantitative analysis of HLA-G expression intensity in CT30 EVTs obtained by exposure to a S1PR3 agonist, CYM5541 (n=2736) or CYM5541 in the presence of a PKCα/β inhibitor, Gö6976 (n=1633). Analysis was performed in MATLAB and at least 2 biological replicates were used. (***p<0.0005). Data for CYM5541 is same as used in Figure 3. (E) Schematic of protocol for hTSC differentiation to EVT obtained by exposure to laminin in the presence of a S1PR3 antagonist, TY-51256. (F) Confocal image of EVT obtained by exposure to laminin in the presence of a S1PR3 antagonist, TY-51256 from CT30 hTSCs, staining for HLA-G and Notch1. Nuclei were stained with DAPI. (G) Quantitative analysis of Notch1 expression intensity in CT30 EVTs obtained by exposure to laminin (n=2197) or laminin in the presence of a S1PR3 antagonist, TY-51256 (n=2564). Analysis was performed in MATLAB and at least 2 biological replicates were used. (***p<0.0005). Data for laminin is same as used in Figure 4. (H) Quantitative analysis of HLA-G expression intensity in CT30 EVTs obtained by exposure to laminin (n=2197) or laminin in the presence of a S1PR3 antagonist inhibitor, TY-51256 (n=2564). Analysis was performed in MATLAB and at least 2 biological replicates were used. (ns, not significant). Data for laminin is same as in Figures 3 and 4. Scale bars are 100µm for all images.

To determine if S1PR3 signaling was necessary for laminin-induced EVT differentiation, we conducted EVT differentiation with laminin-1 in the presence of the S1PR3 antagonist, TY-52158 (**Fig. 5E**). We observed significant cell death under these conditions. However, EVT differentiation could be carried out when 4x cells were seeded during initiation of differentiation, and Notch1^+^ colonies and single, HLA-G^+^ mesenchymal cells were observed (**Figs. 5F, S5G, H**). Interestingly, quantitative image analysis showed that expression intensity of Notch1 was consistently higher in the presence of the S1PR3 antagonist across three cell lines (**Figs. 5G, S5I, K**); however, HLA-G expression remained unaffected or increased marginally in presence of the S1PR3 antagonist (**Figs. 5H, S5J, L**) These results showed that although activation of S1PR3 could induce EVT differentiation in the absence of laminin, S1PR3 activation is not necessary for EVT differentiation mediated by laminin exposure.

## Discussion

Human trophoblast stem cells derived from the trophectoderm layer of blastocyst-stage embryos, first trimester placentas, or human pluripotent stem cells can model the CTB cells during early placental development in vivo. However, the use of forskolin during STB differentiation of hTSCs or the presence of a TGFβ inhibitor and a passage step during EVT differentiation impede mechanistic studies on hTSC differentiation in vitro. Here we present chemically defined conditions for hTSC differentiation to STB and EVT in vitro. We show that hTSCs differentiate to STB over a 6 day period, in the absence of forskolin, in a chemically defined medium that is supplemented with EGF and a ROCK inhibitor. Strikingly, short term (2 days) exposure to a single additional factor during early differentiation, laminin-1, switches the terminal differentiation fate of hTSCs to the EVT lineage (**Fig. 6**). Notably, differentiation to EVT under these conditions does not involve TGFβ inhibition or an intermediate passage step.

**Figure 6:**
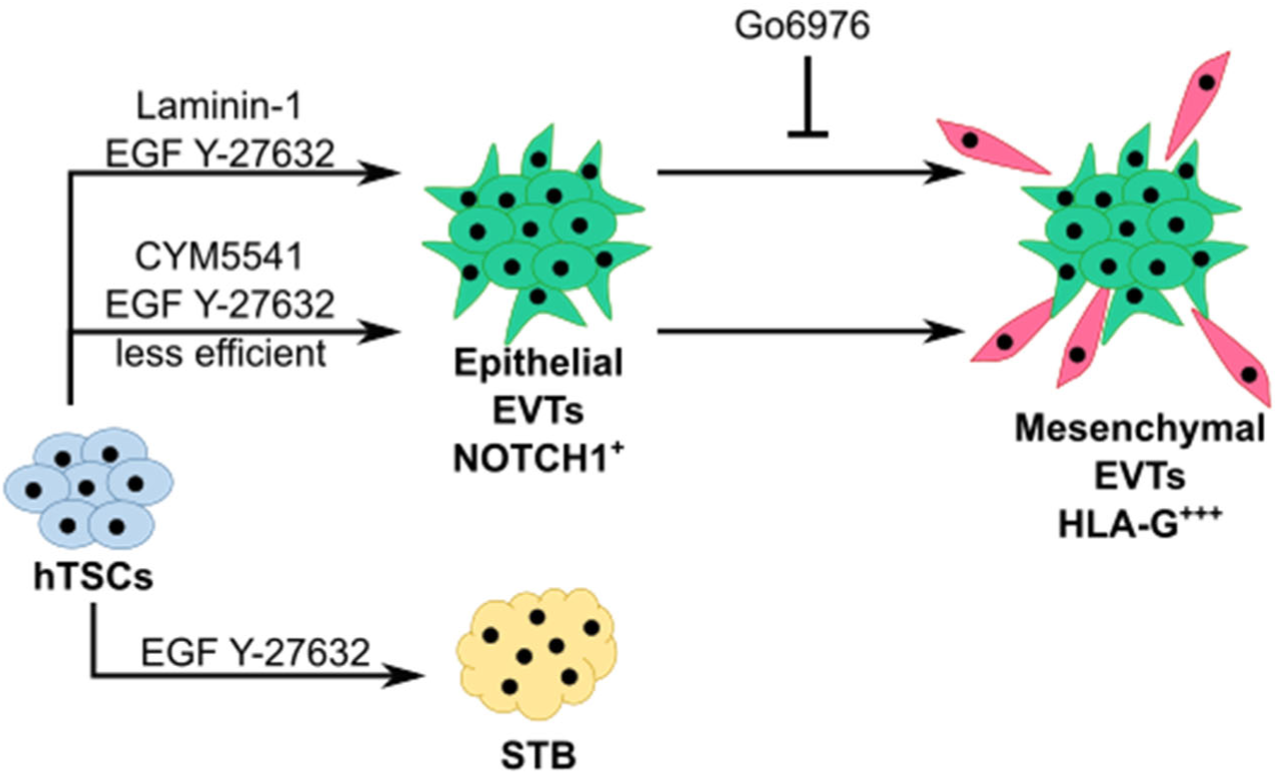
Laminin-1 switches the terminal trophoblast differentiation fate from STB to EVT. Schematic of protocol for hTSC differentiation to STB and EVT obtained by exposure to a laminin or the S1PR3 agonist, CYM5541.

### Cues from extracellular matrix

Exposure to laminin acts as a critical extracellular cue to direct hTSC differentiation to the EVT lineage in vitro. Importantly, a high concentration of laminin is used in our protocol, resulting in the formation of thin layer of substrate overlaying the cells in culture (**Fig. S2A**). Our results are consistent with previous studies by Okae et al., where soluble Matrigel™ was added to cell culture during the first step of differentiation in a two-step protocol^31^. Further, our results are also consistent with previous studies on EVT differentiation of trophoblast derived from human embryonic stem cells (hESCs)^39^. hESC-derived trophoblast underwent differentiation to STB, but not EVT, in the absence of TGFβ inhibition in 2D culture; however, EVT differentiation was obtained in a 3D culture with Matrigel™ with the same culture medium. Taken together with these previous studies, our results implicate a possible role for cues from the extracellular matrix during EVT differentiation. Previous studies that report loss of invasion in primary first trimester CTBs upon treatment with antibodies against laminin further underscore the role of signals from laminin in EVT differentiation^73^. Nevertheless, it is important to note that signaling downstream of multiple extracellular cues likely contributes to EVT differentiation. Indeed, our results show that EVT differentiation of hTSCs could occur in the absence of laminin exposure when S1PR3 was activated using a small molecule agonist.

### A single-step protocol captures heterogeneity of cell types and illustrates the multi-step process of hTSC differentiation to EVTs

A notable feature of our EVT differentiation protocol is the absence of a passage step that has been previously used to form mesenchymal HLA-G^+^ EVTs^31^. The absence of a passage step enables the use of quantitative image analysis to capture of the heterogeneity of cell types that arise as epithelial hTSCs differentiate to mature mesenchymal EVTs. Our results show that expression of Notch1 is upregulated early during EVT differentiation, but downregulated as HLA-G expression increases and mesenchymal EVTs are formed. These results are consistent with observations in vivo, where Notch1 expression is higher in proximal column trophoblasts, and decreases in the distal column where EMT occurs ^3,52,54,57,64,65^. Overall, our results suggest that hTSC differentiation in vitro can be considered as a two-step process with an initial increase in Notch1 expression in epithelial-like cells, followed by downregulation of Notch1 and increased HLA-G expression in mature mesenchymal cells (**Fig. 6**).

### Role of PKCα/β signaling in EVT differentiation

Our results show that inhibition of PKCα/β signaling during laminin exposure resulted in a significant decrease in HLA-G expression; a near complete absence of HLA-G^+^ mesenchymal cells was observed during immunofluorescence analysis. Concomitantly, we observed an increase in the expression of Notch1, as well as increased expression levels of undifferentiated CTB markers (**Figs. 4C-E, S4C-E**). On the other hand, inhibition of PKCα/β signaling does not affect STB differentiation. Taken together these results suggest that PKCα/β signaling is important for EVT differentiation to mature HLA-G^+^ mesenchymal EVTs; however, inhibition of PKCα/β does not prevent commitment to the EVT lineage during hTSC differentiation. Interestingly, the effects of PKCα/β inhibition on EVT differentiation find striking parallels in the cancer literature. PKCα attenuates Notch1 signaling in ErbB2-positive breast cancer cells; inhibition of PKCα increases Notch1 signaling^74^. Inhibition of PKCα also suppresses EMT in breast cancer stem cells, similar to significant reduction in HLA-G^+^ mesenchymal EVTs when PKCα/β signaling is inhibited during hTSC differentiation^75^. However, in contrast to its effects on EVT differentiation mediated by laminin exposure, PKCα/β inhibition did not significantly affect EVT differentiation mediated by S1PR3 activation. Overall, HLA-G expression was lower during EVT differentiation due to S1PR3 activation in the absence of laminin relative to laminin exposure. Nevertheless, HLA-G expression intensity or the formation of mesenchymal EVTs was unaffected by PKCα/β inhibition during S1PR3 agonist-mediated EVT differentiation. Further, laminin-mediated EVT differentiation could occur in the presence of S1PR3 inhibition, although significant reduction in cell number was observed. Taken together, these results suggest that laminin and S1PR3 activation can mediate EVT differentiation through intracellular pathways that may be at least partially non-overlapping.

In conclusion, we have described a chemically defined culture system for hTSC differentiation in vitro that overcome limitations of current approaches. Our results provide baseline conditions for future mechanistic studies on hTSC differentiation.

## Methods

### Key resources

Key resources for this study are listed in **Table 1**.

**Table 1.**
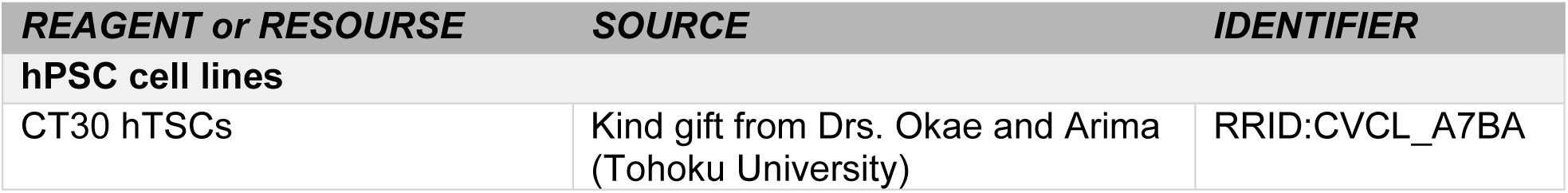
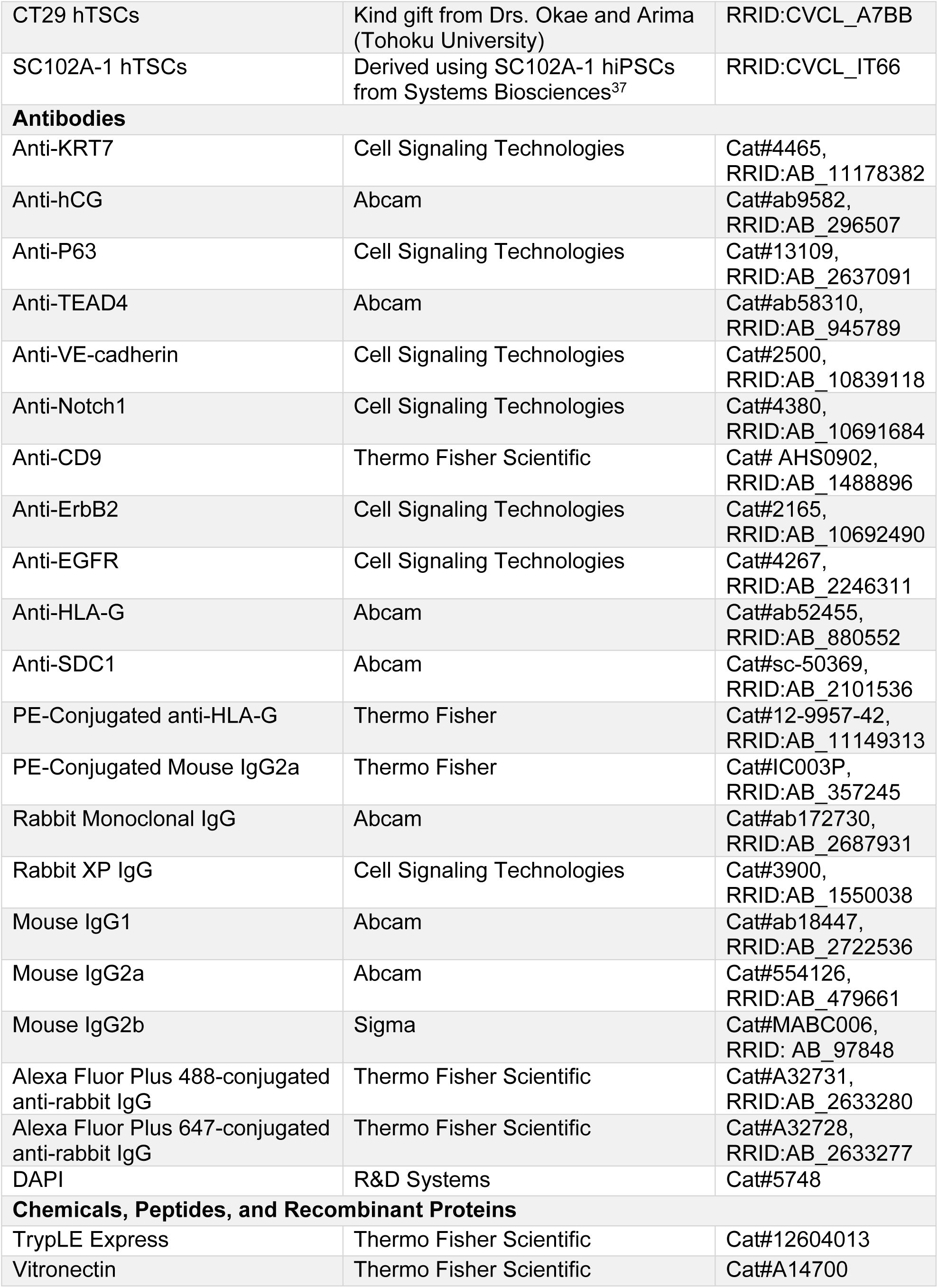
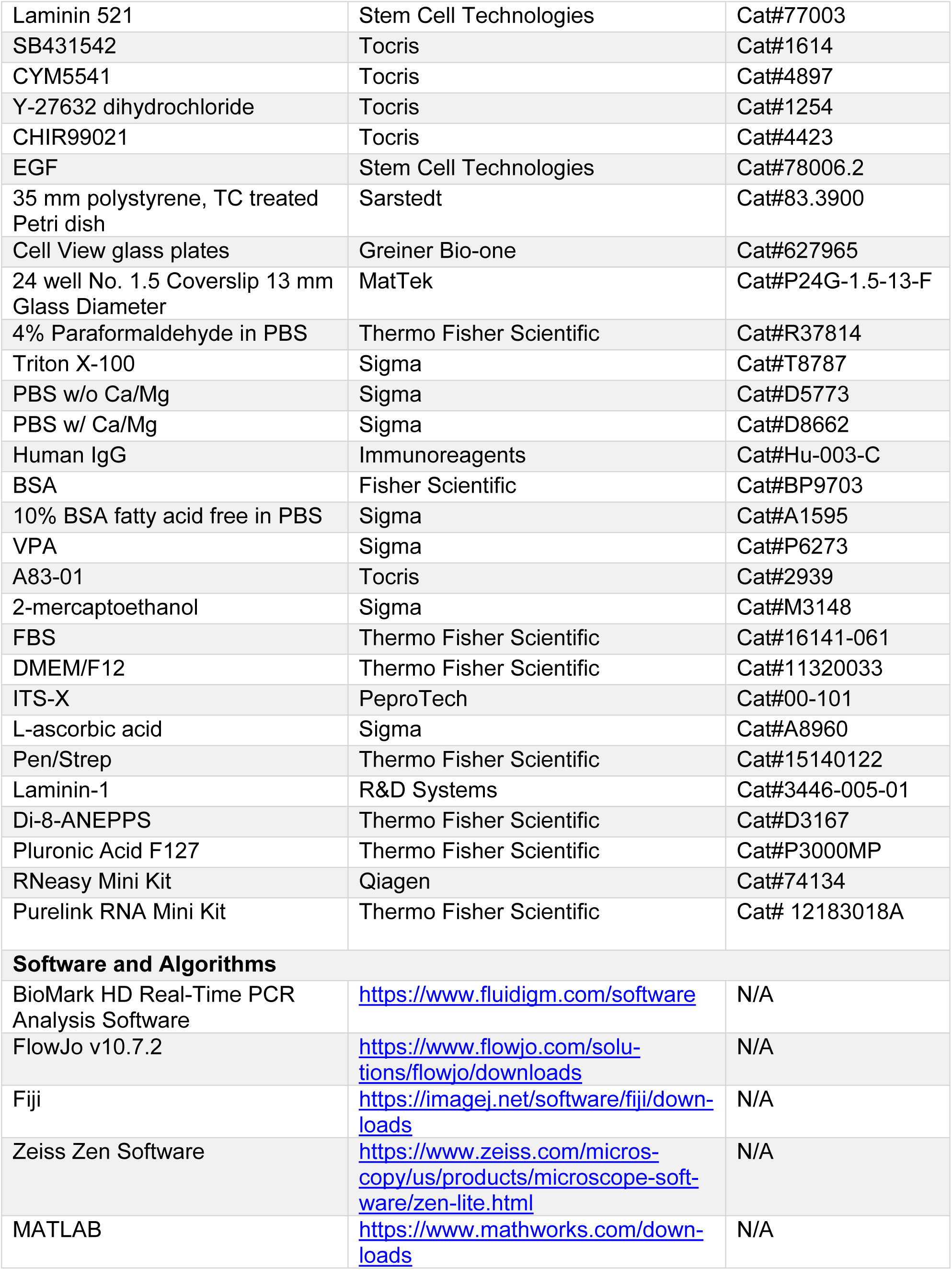
Key Resources.

#### hTSC cell culture

hTSCs were cultured as previously described (Okae et al., 2018) with minor modifications. Cells were cultured in 2 mL of TSCM medium [Dulbecco’s Modified Eagle Medium/Nutrient Mixture F-12 (DMEM/F-12) supplemented with 0.1 mM 2-mercaptoethanol, 0.2% FBS, 0.5% Penicillin-Streptomycin (Pen/Strep), 0.3% BSA, 1% Insulin-Transferrin-Selenium-Ethanolamine (ITS-X), 1.5 µg/mL L-ascorbic acid, 50 ng/mL EGF, 2 μM CHIR99021, 0.5 μM A83-01, 1 μM SB431542, 0.8 mM VPA, and 5 μM Y-27632] at 37°C and 5% CO_2,_ on 35 mm polystyrene plates, pre-coated with 3 µg/ml of vitronectin and 1 µg/ml of Laminin 521. Culture medium was replaced every two days. When cells reached confluence, they were dissociated with TrypLE Express for 10-15 minutes at 37°C and passaged at a 1:10 split ratio. Cells were routinely passaged approximately every 4-6 days. All hTSCs used in this study were passaged at least 5 times prior to use in experiments.

#### EVT and STB differentiation

Prior to differentiation, hTSCs at confluence were dissociated into single cells using TrypLE Express and 1.5×10^5^ cells were seeded onto a new 35 mm polystyrene plate pre-coated plate with 3 µg/ml of vitronectin and 1 µg/ml of Laminin 521. For STB differentiation, cells were cultured in defined trophoblast differentiation medium (DTDM) [DMEM/F-12 supplemented with 1% ITS-X, 75 µg/mL L-ascorbic acid]. For EVT differentiation, DTDM was supplemented with 150 µg/mL laminin-1 after cells were plated in DTDM. 5 µM Y-27632 and 50 ng/mL EGF was added at passage. Cell culture medium was replaced every 2 days and cultures were analyzed at day 6 unless otherwise specified.

#### Immunostaining

For immunofluorescence analysis, 3×10^4^ cells were grown on 24-well glass bottom plates coated with 3 µg/ml of vitronectin and 1 µg/ml of Laminin 521. Where indicated, 2x or 4x this number of cells were plated. Cells were fixed with 4% paraformaldehyde fixative solution for 5 minutes, permeabilized with 0.5% Triton X-100 in PBS for 10 minutes, then blocked in blocking buffer [0.5% BSA, and 200 μM human IgG in PBS] for at least one hour. Cells were then incubated overnight at 4°C in primary antibody diluted in blocking buffer. Primary antibodies used were anti-HLA-G (1:250), anti-VE-cadherin (1:250), and anti-Notch1 (1:200), anti-CD9 (1:50), anti-ErbB2 (1:250), anti-EGFR (1:50), anti-hCG (1:50), anti-SDC-1 (1:250), anti-KRT7 (1:50), anti-p63 (1:250). Secondary antibodies were added an hour before imaging. Corresponding isotype controls (rabbit monoclonal IgG, rabbit XP IgG, mouse IgG1, mouse IgG2a, and mouse IgG2b) were used at primary antibody concentrations. Alexa Fluor 488- or Alexa Fluor 647-conjugated secondary antibodies were used. Nuclei were stained with DAPI and images were taken with a laser scanning confocal microscope (LSM880, Carl Zeiss, Germany).

#### Confocal Image Analysis

Image analysis was conducted using an image processing algorithm created in MATLAB R2018a. All image processing was performed *post hoc*. First, DAPI (blue channel) was isolated from the RBG image, binarized, and processed to accurately represent the number of cells in each image. The primary-antibody stain of interest was isolated and processed in the same manner. This was performed for one control image and multiple experimental images. Each cell in the experimental images was considered positively stained if the average intensity of that cell was greater than the average intensity of all of the cells in the control image. Statistical analysis was conducted using a two-tailed t-test.

#### Membrane staining

Differentiated cells were fixed, permeabilized, and blocked as previously described. 2.5 µL of pluronic acid was added to 25 µM Di-8-ANEPPS in PBS and allowed to incubate for one hour at 4 °C. For the nuclear stain, 1 μL of 4, 6-diamidino-2-phenylindole (DAPI) was added to each plate and incubated for 5 – 10 minutes. Pluronic acid, Di-8-ANEPPS and DAPI were removed by washing twice with PBS with Ca/Mg before imaging using a Keyence BZ-X810 system.

#### Flow Cytometry

Cells were dissociated with TrypLE and fixed with 2% paraformaldehyde for 10 minutes in suspension. They were then blocked in Saponin blocking buffer [1% BSA 1 mg/mL Saponin] for 15 minutes at room temperature. PE-conjugated anti-HLA-G antibody (1:20) or anti-Notch1 antibody (1:800) diluted in Saponin blocking buffer was then added and the cells incubated for 1 hour on ice. Alexa-Fluor Plus 488-conjugated rabbit IgG was added as a secondary to Notch1 antibody staining and incubated for an additional hour on ice. PE-conjugated mouse IgG2a or Rabbit XP IgG was used as the isotype control. Flow cytometry was carried out using a MACSQuant VYB, and the data were analyzed using FlowJo software.

#### RNA extraction, cDNA synthesis and qRT-PCR

RNA was isolated using Invitrogen PureLink RNA Mini Kit (Fisher Scientific) or the RNeasy Mini Kit (Qiagen) using the manufacturers’ protocol. RNA was quantified using a Nanodrop 1000 spectrophotometer (Thermo Scientific, Waltham, MA). 1 μL of RNA was transformed into complementary DNA using Fluidigm Reverse Transcription Master Mix (Fluidigm) according to the manufacturer’s protocol and underwent 16 preamplification cycles using the Fluidigm Preamp Master Mix according to the manufacturer’s protocol. RNA was then analyzed by the UNC Advanced Analytics Core facility using the Fluidigm Biomark HD 96.96 IFC array (Fluidigm) and validated TaqMan probes according to the manufacturer’s protocol. Using BioMark HD software (Fluidigm), Ct values were then normalized against the geometric mean of glyceraldehyde 3-phosphate dehydrogenase (GAPDH) and beta-actin (ACTB), and relative log_2_fold changes were calculated, normalizing to day 0 hTSCs based on the ΔΔCT method (Livak and Schmittgen, 2001). Statistical analysis was conducted using a two-tailed t-test with unequal variance.

## Author contributions

Conceptualization: VK, AM, BR; Investigation: VK, TM, AC; Formal Analysis: VK, AC, BR; Resources: ASM; Writing: VK, BR; Visualization: VK, BR; Project Administration: BR; Funding Acquisition: BR and ASM

## Acknowledgments

This work was supported by NIH grants HD092741, HD093982 and NSF grant CBET 1706118.

## Conflict of interest

The authors declare that they have no conflicts of interest with the contents of this article.

## Supplementary Information

Supplementary Information.pdf (contains Supplementary Figures S1-S6)

Tables S1-S2.xlsx (contains Supplementary Tables S1, S2)

## Notes

### Competing Interest Statement

The authors have declared no competing interest.

## References

1. Aplin, J. D. Developmental cell biology of human villous trophoblast: Current research problems. International Journal of Developmental Biology 54, 323–329 (2010).

2. Knöfler, M., Vasicek, R. & Schreiber, M. Key regulatory transcription factors involved in placental trophoblast development - A review. Placenta 22, (2001).

3. Knöfler, M. et al. Human placenta and trophoblast development: key molecular mechanisms and model systems. Cellular and Molecular Life Sciences vol. 76 3479–3496 (2019).

4. Loregger, T., Pollheimer, J. & Knöfler, M. Regulatory transcription factors controlling function and differentiation of human trophoblast - A review. Placenta 24, (2003).

5. Caniggia, I., Winter, J., Lye, S. J. & Post, M. Oxygen and placental development during the first trimester: Implications for the pathophysiology of pre-eclampsia. Placenta 21, (2000).

6. Tuuli, M. G., Longtine, M. S. & Nelson, D. M. Review: Oxygen and trophoblast biology - A source of controversy. in Placenta vol. 32 (2011).

7. Zhou, Y., Genbacev, O., Damsky, C. H. & Fisher, S. J. Oxygen regulates human cytotrophoblast differentiation and invasion: Implications for endovascular invasion in normal pregnancy and in pre-eclampsia. in Journal of Reproductive Immunology vol. 39 197–213 (1998).

8. Vićovac, L. & Aplin, J. D. Epithelial-mesenchymal transition during trophoblast differentiation. Cells Tissues Organs 156, 202–216 (1996).

9. Thiery, J.P. Epithelial-mesenchymal transitions in development and pathologies. Current Opinion in Cell Biology vol. 15 740–746 (2003).

10. Dasilva-Arnold, S., James, J. L., Al-Khan, A., Zamudio, S. & Illsley, N. P. Differentiation of first trimester cytotrophoblast to extravillous trophoblast involves an epithelial-mesenchymal transition. Placenta 36, 1412–1418 (2015).

11. Benirschke, Kurt., Burton, G. (Graham J.) & Baergen, R. N. Pathology of the human placenta. (Springer, 2012).

12. Soares, M. J., Iqbal, K. & Kozai, K. Hypoxia and Placental Development. Birth Defects Research vol. 109 1309–1329 (2017).

13. Harris, L. K. et al. Invasive trophoblasts stimulate vascular smooth muscle cell apoptosis by a Fas ligand-dependent mechanism. American Journal of Pathology 169, 1863–1874 (2006).

14. Lash, G. E. et al. Decidual macrophages: key regulators of vascular remodeling in human pregnancy. Journal of Leukocyte Biology 100, 315–325 (2016).

15. Pijnenborg, R., Vercruysse, L. & Hanssens, M. The Uterine Spiral Arteries In Human Pregnancy: Facts and Controversies. Placenta vol. 27 939–958 (2006).

16. Raymond, D. & Peterson, E. A critical review of early-onset and late-onset preeclampsia. Obstetrical and Gynecological Survey vol. 66 497–506 (2011).

17. Prossler, J., Chen, Q., Chamley, L. & James, J. L. The relationship between TGFβ, low oxygen and the outgrowth of extravillous trophoblasts from anchoring villi during the first trimester of pregnancy. Cytokine 68, 9–15 (2014).

18. Arnholdt, H., Meisel, F., Fandrey, K. & Löhrs, U. Proliferation of villous trophoblast of the human placenta in normal and abnormal pregnancies. Virchows Archiv B Cell Pathology Including Molecular Pathology 60, 365–372 (1991).

19. Silver, R. M. Examining the link between placental pathology, growth restriction, and stillbirth. Best Practice and Research: Clinical Obstetrics and Gynaecology vol. 49 89–102 (2018).

20. Naicker, T., Khedun, S. M., Moodley, J. & Pijnenborg, R. Quantitative analysis of trophoblast invasion in preeclampsia. Acta Obstetricia et Gynecologica Scandinavica 82, 722–729 (2003).

21. Zhou, Y., Damsky, C. H. & Fisher, S. J. Preeclampsia is associated with failure of human cytotrophoblasts to mimic a vascular adhesion phenotype: One cause of defective endovascular invasion in this syndrome? Journal of Clinical Investigation 99, 2152–2164 (1997).

22. Zhou, Y. et al. Reversal of gene dysregulation in cultured cytotrophoblasts reveals possible causes of preeclampsia. 123, 2862–2872 (2013).

23. Boer, G. J. Ethical guidelines for the use of human embryonic or fetal tissue for experimental and clinical neurotransplantation and research. Journal of Neurology 242, 1–13 (1994).

24. Pera, M. F. Human embryo research and the 14-day rule. Development (Cambridge) 144, 1923–1925 (2017).

25. Soncin, F. et al. Comparative analysis of mouse and human placentae across gestation reveals species-specific regulators of placental development. Development (Cambridge) 145, (2018).

26. Lee, K. Y. & Demayo, F. J. Animal models of implantation. Reproduction 128, 679–695 (2004).

27. Malassine, A., Frendo, J.-L. & Evain-Brion, D. A comparison of placental development and endocrine functions between the human and mouse model. doi:10.1093/humupd/dmg043.

28. Kunath, T. et al. Developmental differences in the expression of FGF receptors between human and mouse embryos. Placenta 35, 1079–1088 (2014).

29. Bilban, M. et al. Trophoblast invasion: Assessment of cellular models using gene expression signatures. Placenta 31, 989–996 (2010).

30. Hannan, N. J., Paiva, P., Dimitriadis, E. & Salamonsen, L. A. Models for Study of Human Embryo Implantation: Choice of Cell Lines?1. Biology of Reproduction 82, 235–245 (2010).

31. Okae, H. et al. Derivation of Human Trophoblast Stem Cells. Cell Stem Cell 22, 50–63.e6 (2018).

32. Yu, L. et al. Blastocyst-like structures generated from human pluripotent stem cells. Nature 2021 591:7851 591, 620–626 (2021).

33. Amita, M. et al. Complete and unidirectional conversion of human embryonic stem cells to trophoblast by BMP4. Proceedings of the National Academy of Sciences of the United States of America 110, (2013).

34. Castel, G. et al. Induction of Human Trophoblast Stem Cells from Somatic Cells and Pluripotent Stem Cells. Cell Reports 33, 108419 (2020).

35. Wei, Y. et al. Efficient derivation of human trophoblast stem cells from primed pluripotent stem cells. Science Advances 7, eabf4416 (2021).

36. Dong, C. et al. Derivation of trophoblast stem cells from naïve human pluripotent stem cells. ELife 9, (2020).

37. Mischler, A. et al. Two distinct trophectoderm lineage stem cells from human pluripotent stem cells. Journal of Biological Chemistry 296, (2021).

38. Tiruthani, K., Sarkar, P. & Rao, B. Trophoblast differentiation of human embryonic stem cells. Biotechnology Journal 8, 421–433 (2013).

39. Sarkar, P. et al. Activin/nodal signaling switches the terminal fate of human embryonic stem cell-derived trophoblasts. Journal of Biological Chemistry 290, 8834–8848 (2015).

40. Cinkornpumin, J. K. et al. Naive Human Embryonic Stem Cells Can Give Rise to Cells with a Trophoblast-like Transcriptome and Methylome. Stem Cell Reports 15, 198–213 (2020).

41. Sheridan, M. A. et al. Early onset preeclampsia in a model for human placental trophoblast. Proceedings of the National Academy of Sciences 116, 4336–4345 (2019).

42. Caniggia, I., Grisaru-Gravnosky, S., Kuliszewsky, M., Post, M. & Lye, S. J. Inhibition of TGF-β3 restores the invasive capability of extravillous trophoblasts in preeclamptic pregnancies. Journal of Clinical Investigation 103, 1641–1650 (1999).

43. Lala, P. K. & Graham, C. H. Mechanisms of trophoblast invasiveness and their control: the role of proteases and protease inhibitors. CANCER AND METASTASIS REVIEW 9, 369–379 (1990).

44. Cheng, J. C., Chang, H. M. & Leung, P. C. K. Transforming growth factor-β1 inhibits trophoblast cell invasion by inducing snail-mediated down-regulation of vascular endothelial-cadherin protein. Journal of Biological Chemistry 288, 33181–33192 (2013).

45. Pollheimer, J. & Knöfler, M. Signalling pathways regulating the invasive differentiation of human trophoblasts: A review. Placenta 26, S21–S30 (2005).

46. Xu, J. et al. Aberrant TGFβ signaling contributes to altered trophoblast differentiation in preeclampsia. Endocrinology 157, 883–899 (2016).

47. Graham, C. H. & Lala, P. K. Mechanism of control of trophoblast invasion in situ. Journal of Cellular Physiology 148, 228–234 (1991).

48. Haider, S., Kunihs, V., Fiala, C., Pollheimer, J. & Knöfler, M. Expression pattern and phosphorylation status of Smad2/3 in different subtypes of human first trimester trophoblast. Placenta 57, 17–25 (2017).

49. Romero, R., Kusanovic, J. P., Chaiworapongsa, T. & Hassan, S. S. Placental bed disorders in preterm labor, preterm PROM, spontaneous abortion and abruptio placentae. Best Practice and Research: Clinical Obstetrics and Gynaecology vol. 25 313–327 (2011).

50. Fryer, B. H. & Simon, M. C. Hypoxia, HIF and the placenta. Cell Cycle vol. 5 495–498 (2006).

51. Rosati, E. et al. Constitutively activated Notch signaling is involved in survival and apoptosis resistance of B-CLL cells. Blood 113, 856–865 (2009).

52. Haider, S. et al. Self-Renewing Trophoblast Organoids Recapitulate the Developmental Program of the Early Human Placenta. Stem Cell Reports 11, 537–551 (2018).

53. Haider, S. et al. Notch signaling plays a critical role in motility and differentiation of human first-trimester cytotrophoblasts. Endocrinology 155, 263–274 (2014).

54. Haider, S. et al. Notch1 controls development of the extravillous trophoblast lineage in the human placenta. Proceedings of the National Academy of Sciences of the United States of America 113, E7710–E7719 (2016).

55. Meinhardt, G. et al. Wnt-Dependent T-Cell Factor-4 controls human etravillous trophoblast motility. Endocrinology 155, 1908–1920 (2014).

56. Hunkapiller, N. M. et al. A role for Notch signaling in trophoblast endovascular invasion and in the pathogenesis of pre-eclampsia. Development 138, 2987–2998 (2011).

57. Plessl, K., Haider, S., Fiala, C., Pollheimer, J. & Knöfler, M. Expression pattern and function of Notch2 in different subtypes of first trimester cytotrophoblast. Placenta 36, 365–371 (2015).

58. Meinhardt, G., Kaltenberger, S., Fiala, C., Knöfler, M. & Pollheimer, J. ERBB2 gene amplification increases during the transition of proximal EGFR+ to distal HLA-G+ first trimester cell column trophoblasts. Placenta 36, 803–808 (2015).

59. Benirschke, K. & Driscoll, S. G. The Pathology of the Human Placenta. in Placenta 97–571 (Springer Berlin Heidelberg, 1967). doi:10.1007/978-3-662-38455-8_2.

60. Pawar, S. C., Demetriou, M. C., Nagle, R. B., Bowden, G. T. & Cress, A. E. Integrin α6 cleavage: A novel modification to modulate cell migration. Experimental Cell Research 313, 1080–1089 (2007).

61. Mccormick, J., Whitley, G. S. J., Le Bouteiller, P. & Cartwright, J. E. Soluble HLA-G regulates motility and invasion of the trophoblast-derived cell line SGHPL-4. 24, 1339–1345 (2009).

62. Fukushima, K. et al. Tumor Necrosis Factor and Vascular Endothelial Growth Factor Induce Endothelial Integrin Repertories, Regulating Endovascular Differentiation and Apoptosis in a Human Extravillous Trophoblast Cell Line1. Biology of Reproduction 73, 172–179 (2005).

63. Aisenbrey, E. A. & Murphy, W. L. Synthetic alternatives to Matrigel. Nature Reviews Materials 2020 5:7 5, 539–551 (2020).

64. Haider, S. et al. Notch signaling plays a critical role in motility and differentiation of human first-trimester cytotrophoblasts. Endocrinology 155, 263–274 (2014).

65. Benirschke, K., Burton, G. J. & Baergen, R. N. Pathology of the human placenta, sixth edition. Pathology of the Human Placenta (Springer Berlin Heidelberg, 2012). doi:10.1007/978-3-642-23941-0.

66. Pathology of the Human Placenta. Pathology of the Human Placenta (Springer New York, 2006). doi:10.1007/b137920.

67. Wang, J., Mayernik, L. & Armant, D. R. Integrin Signaling Regulates Blastocyst Adhesion to Fibronectin at Implantation: Intracellular Calcium Transients and Vesicle Trafficking in Primary Trophoblast Cells. Developmental Biology 245, 270–279 (2002).

68. Randall, D. Integrin-Mediated Adhesion and Signaling during Blastocyst Implantation. Cells Tissues Organs 172, 3 (2002).

69. Taylor, J., Pampillo, M., Bhattacharya, M. & Babwah, A. v. Kisspeptin/KISS1R signaling potentiates extravillous trophoblast adhesion to type-I collagen in a PKC- and ERK1/2-dependent manner. Molecular Reproduction and Development 81, 42–54 (2014).

70. Fakhr, Y., Brindley, D. N. & Hemmings, D. G. Physiological and pathological functions of sphingolipids in pregnancy. Cellular Signalling 85, 110041 (2021).

71. Gandy, K. A. O. & Obeid, L. M. Regulation of the Sphingosine Kinase/Sphingosine 1-Phosphate Pathway. 275–303 (2013) doi:10.1007/978-3-7091-1511-4_14.

72. Rennecke, J. et al. Immunological Demonstration of Protein Kinase Cμ in Murine Tissues and Various Cell Lines. European Journal of Biochemistry 242, 428–432 (1996).

73. Damsky, C. H. et al. Integrin switching regulates normal trophoblast invasion. Development 120, 3657–3666 (1994).

74. Pandya, K. et al. PKCα Attenuates Jagged-1–Mediated Notch Signaling in ErbB-2–Positive Breast Cancer to Reverse Trastuzumab Resistance. Clinical Cancer Research 22, 175–186 (2016).

75. Tam, W. L. et al. Protein Kinase C α Is a Central Signaling Node and Therapeutic Target for Breast Cancer Stem Cells. Cancer Cell 24, 347–364 (2013).

