## Supporting Information for "Laminin switches terminal differentiation fate of human trophoblast stem cells under chemically defined culture conditions"

Supplementary Figures S1-S6

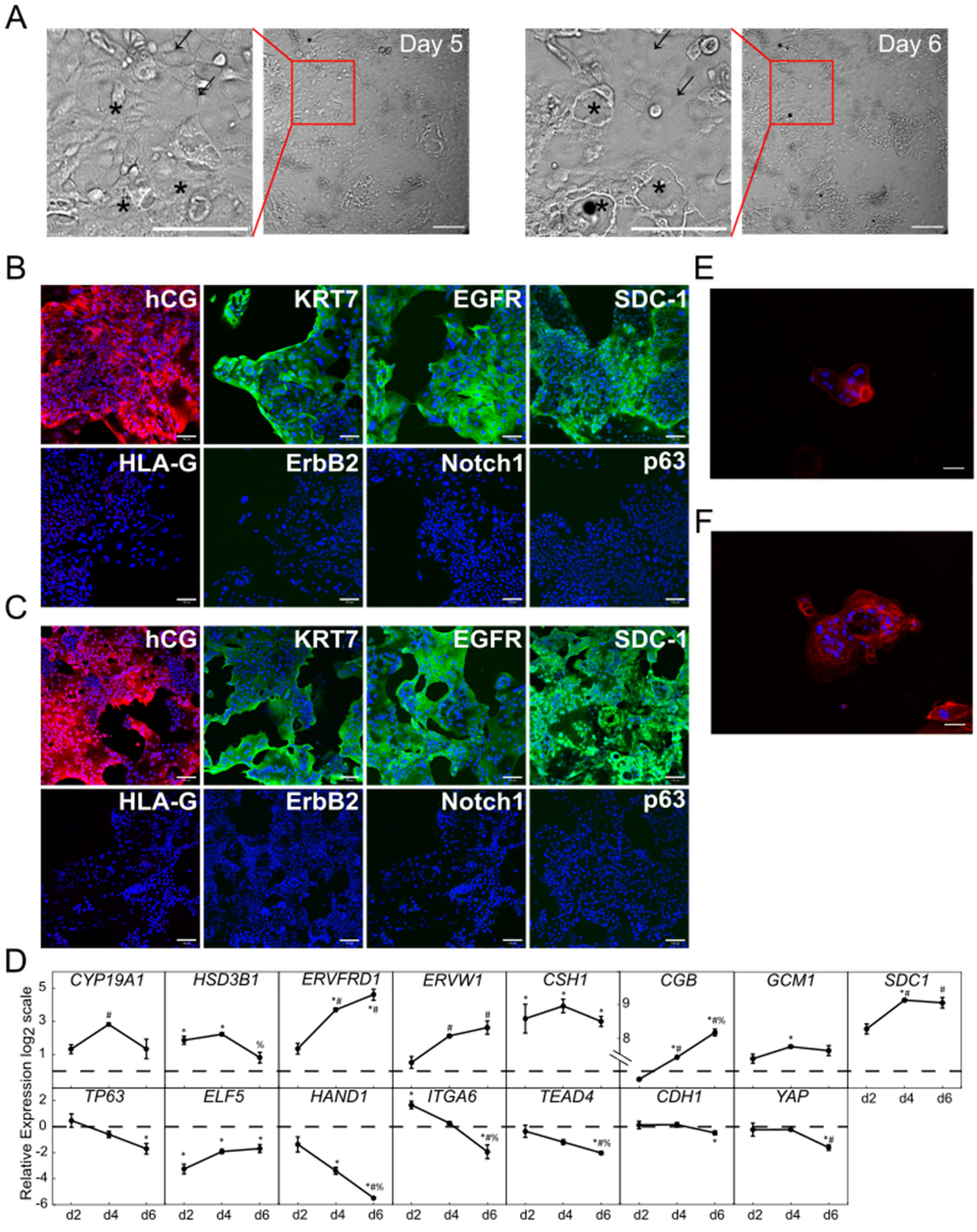

A
Day 5
Day 6
B
hCG
KRT7
EGFR
SDC-1
HLA-G
ErbB2
Notch1
p63
C
hCG
KRT7
EGFR
SDC-1
HLA-G
ErbB2
Notch1
p63
E
F
D
CYP19A1
HSD3B1
ERVFRD1
ERVW1
CSH1
CGB
GCM1
SDC1
TP63
ELF5
HAND1
ITGA6
TEAD4
CDH1
YAP
Relative Expression log2 scale
d2
d4
d6

**Figure S1: Chemically defined conditions for STB differentiation in the absence of forskolin.**

(A) Bright field images of CT30 hTSC differentiation to STB on day 5 and day 6. Outcrop is the magnified image. On day 6, stars indicate where lacunae formed, and arrows indicate where cell boundaries fused. In these same positions on day 5, no lacunae are seen, and distinct cell boundaries are present.

(B) Confocal images of STB from CT29 hTSCs, staining for hCG, KRT7, EGFR, SDC-1, HLA-G, ErbB2, Notch1, p63. Nuclei were stained with DAPI.

(C) Confocal images of STB from SC102A-1 hTSCs, staining for hCG, KRT7, EGFR, SDC-1, HLA-G, ErbB2, Notch1, p63. Nuclei were stained with DAPI.

(D) Gene expression of *CYP19A1, HSD3B1, ERVFRD1, ERVW1, CSH1, CGB, GCM1, SDC-1, TP63, ELF5, HAND1, ITGA6, TEAD4, CDH1,* and *YAP* of STB from CT29 hTSCs. Three biological replicates were used. (Error bars, S.E., $^{*}p<0.05$ for comparison with undifferentiated hTSCs (dashed line), $^{\#}p<0.05$ for comparison with cells at day 2, $^{\%}p<0.05$ for comparison with cells at day 4).

(E) Fluorescent image of STB from CT29 hTSCs. Nuclei were stained with DAPI. Membrane was stained with Di-8-ANEPPS cell membrane stain.

(F) Fluorescent image of STB from SC102A-1 hTSCs. Nuclei were stained with DAPI. Membrane was stained with Di-8-ANEPPS cell membrane stain.

Scale bars are 100μm for all images.

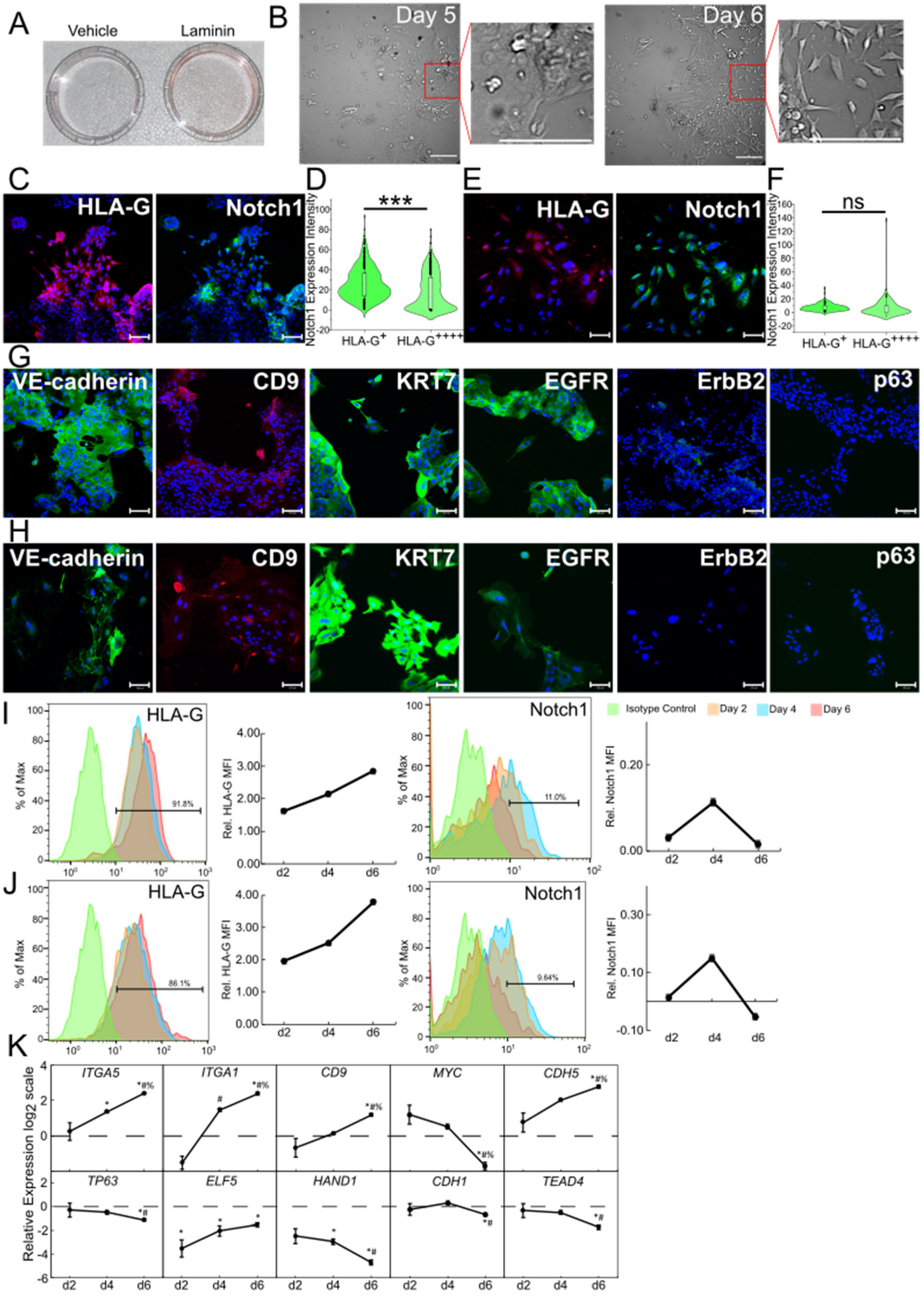
A
Vehicle
Laminin
B
Day 5
Day 6
C
HLA-G
Notch1
D
***
Notch1 Expression Intensity
HLA-G+
HLA-G++++
E
HLA-G
Notch1
F
ns
Notch1 Expression Intensity
HLA-G+
HLA-G++++
G
VE-cadherin
CD9
KRT7
EGFR
ErbB2
p63
H
VE-cadherin
CD9
KRT7
EGFR
ErbB2
p63
I
HLA-G
% of Max
91.8%
Rel. HLA-G MFI
d2
d4
d6
Notch1
% of Max
11.0%
Isotype Control
Day 2
Day 4
Day 6
Rel. Notch1 MFI
J
HLA-G
% of Max
86.1%
Rel. HLA-G MFI
Notch1
9.64%
Rel. Notch1 MFI
K
Relative Expression log2 scale
ITGA5
ITGA1
CD9
MYC
CDH5
TP63
ELF5
HAND1
CDH1
TEAD4
d2
d4
d6

**Figure S2: Presence of laminin switches hTSC differentiation from STB to EVT fate.**

(A) Image of plastic plates with dyed-red vehicle (DMSO) or dyed-red laminin. 75 μL of DMSO or laminin was dyed with red food coloring and was added to 2 mL of DMEM on pre-coated plastic plates. The plates were incubated at 37°C and 5% $CO_2$ for 2 days after which the liquid media was aspirated. The DMSO is completely removed from the plate after aspiration (Vehicle) whereas dyed-red laminin remains present on the plate (Laminin) indicating that a thin layer of laminin gel remains on the plate at this concentration that cannot be aspirated.

(B) Bright field images of CT30 hTSC differentiation to EVT on day 5 and day 6. Outcrop is the magnified image. Cells on day 5 are in an epithelial colony but on day 6 single, mesenchymal cells are observed.

(C) Confocal images of EVT from CT29 hTSCs, staining for HLA-G and Notch1. Nuclei were stained with DAPI.

(D) Quantitative analysis of Notch1 expression intensity of CT29 EVTs from the bottom (HLA-$G^+$) and top (HLA-$G^{++++}$) 25% of HLA-G expression intensity EVTs (n=648, each). Analysis was performed in MATLAB and at least 2 biological replicates were used. (***p<0.0005).

(E) Confocal images of EVT from SC102A-1 hTSCs, staining for HLA-G and Notch1. Nuclei were stained with DAPI.

(F) Quantitative analysis of Notch1 expression intensity of SC102A-1 EVTs from the bottom (HLA-$G^+$) and top (HLA-$G^{++++}$) 25% of HLA-G expression intensity EVTs (n=125, each). Analysis was performed in MATLAB and at least 2 biological replicates were used. (ns, not significant).

(G) Confocal images of EVT from CT29 hTSCs, staining for VE-Cadherin, CD9, KRT7, EGFR, ErbB2, and p63. Nuclei were stained with DAPI.

(H) Confocal images of EVT from SC102A-1 hTSCs, staining for VE-Cadherin, CD9, KRT7, EGFR, ErbB2, and p63. Nuclei were stained with DAPI.

(I) Flow cytometry histogram of HLA-G and Notch1 expression of CT29 EVTs on day 2, day 4, and day 6 compared to an isotype control and their relative mean fluorescence intensity (MFI). Relative MFI is calculated as (X-C)/C where X is the experimental MFI and C is the isotype control MFI.

(J) Flow cytometry histogram of HLA-G and Notch1 expression of SC102A-1 EVTs on day 2, day 4, and day 6 compared to an isotype control and their relative mean fluorescence intensity (MFI).

(K) Gene expression of *ITGA5, CD9, ITGA1, MYC, CDH5, TP63, ELF5, HAND1, CDH1,* and *TEAD4* of EVT from CT29 hTSCs. Three biological replicates were used. (Error bars, S.E., *p<0.05 for comparison with undifferentiated hTSCs (dashed line), $^{\#}$p<0.05 for comparison with cells at day 2, $^{\%}$p<0.05 for comparison with cells at day 4).

Scale bars are 100μm for all images.

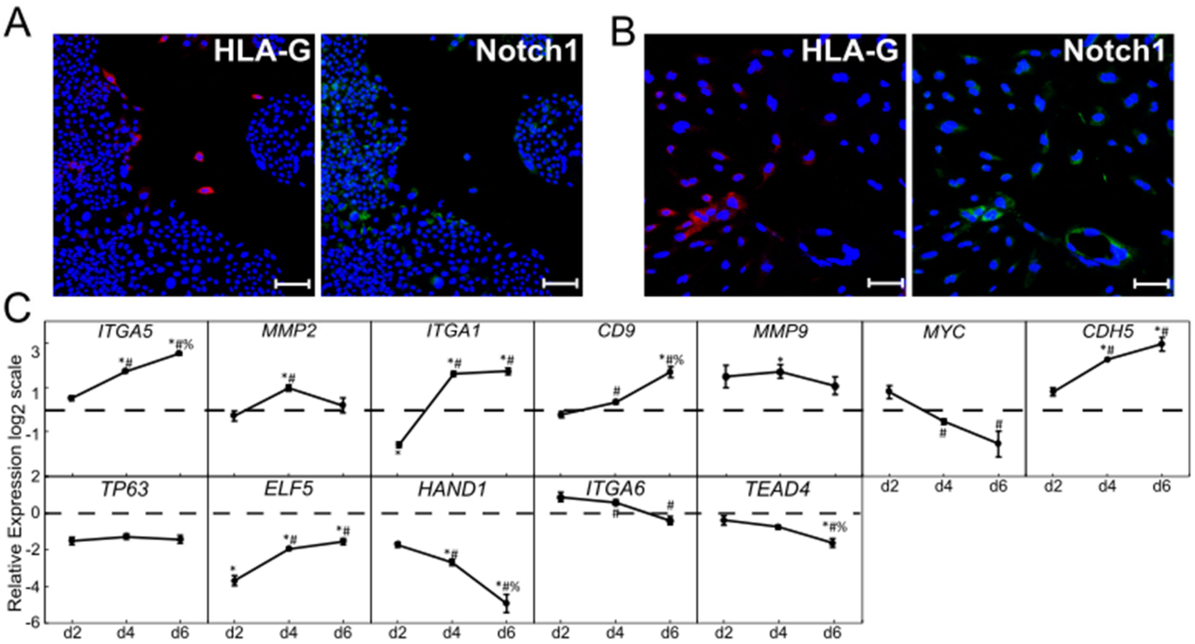


**Figure S3: Activation of S1PR3 agonist can mediate EVT differentiation of hTSCs in the absence of exogenous laminin.**

(A) Confocal images of EVT obtained by exposure to a S1PR3 agonist, CYM5541 from CT29 hTSCs, staining for HLA-G and Notch1. Nuclei were stained with DAPI.

(B) Confocal images of EVT obtained by exposure to a S1PR3 agonist, CYM5541 from SC102A-1 hTSCs, staining for HLA-G and Notch1. Nuclei were stained with DAPI.

(C) Gene expression of *ITGA5, MMP2, ITGA1, CD9, MMP9, MYC, CDH5, TP63, ELF5, HAND1, ITGA6,* and *TEAD4* of EVT obtained by exposure to a S1PR3 agonist, CYM5541 from CT29 hTSCs. Three biological replicates were used. (Error bars, S.E., *$p<0.05$ for comparison with undifferentiated hTSCs (dashed line), $^{\#}p<0.05$ for comparison with cells at day 2, $^{\%}p<0.05$ for comparison with cells at day 4).

Scale bars are 100μm for all images.

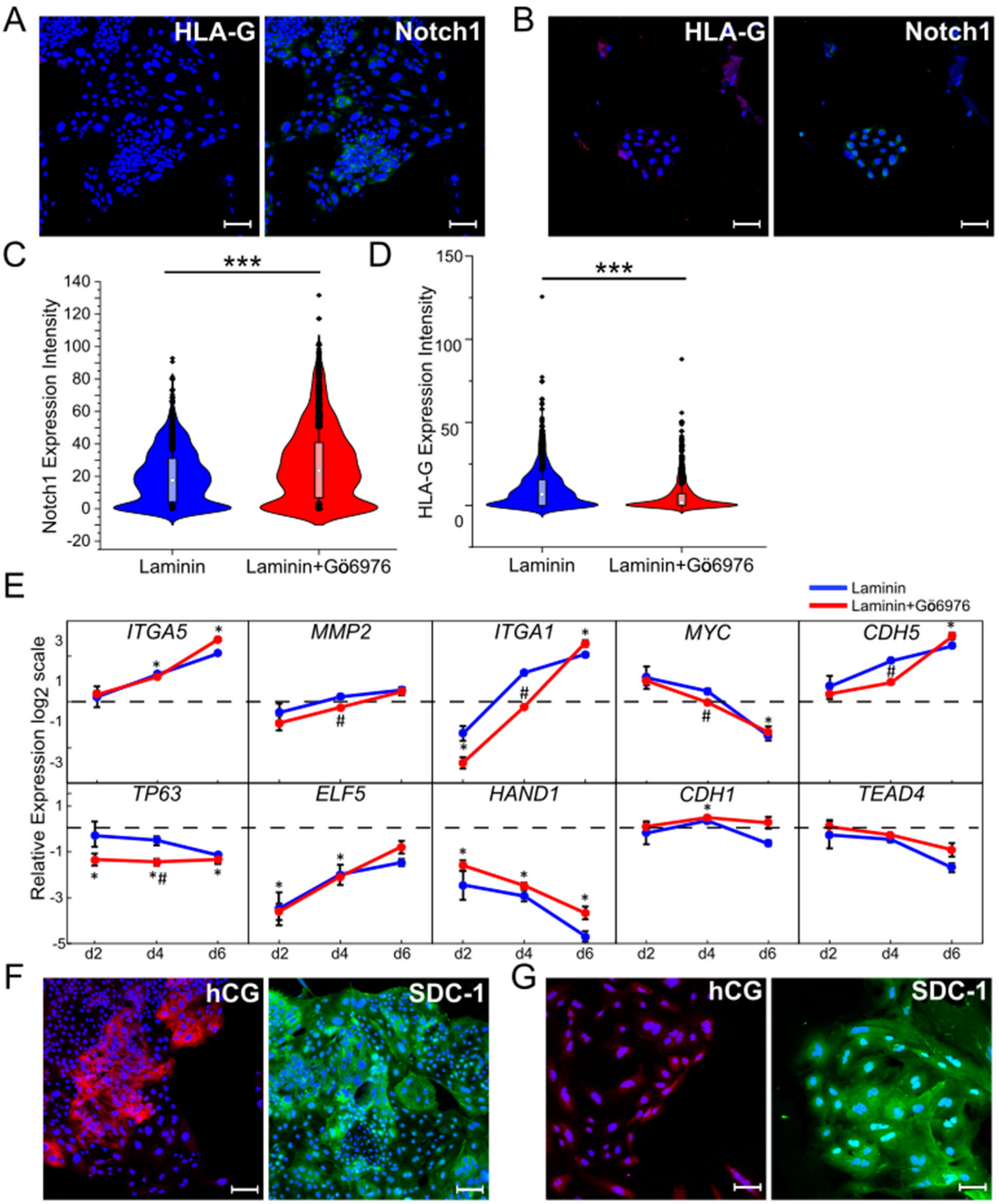


**Figure S4: Inhibition of PKCα/β adversely affects laminin-mediated EVT differentiation.**

(A) Confocal image of EVT obtained by exposure to laminin in the presence of a PKCα/β inhibitor, Gö6976 from CT29 hTSCs, staining for HLA-G and Notch1. Nuclei were stained with DAPI.

(B) Confocal image of EVT obtained by exposure to laminin in the presence of a PKC$\alpha/\beta$ inhibitor, Gö6976 from SC102A-1 hTSCs, staining for HLA-G and Notch1. Nuclei were stained with DAPI.

(C) Quantitative analysis of Notch1 expression intensity in CT29 EVTs obtained by exposure to laminin (n=2592) or laminin in the presence of a PKC$\alpha/\beta$ inhibitor, Gö6976 (n=2436). Analysis was performed in MATLAB and at least 2 biological replicates were used. (***p<0.0005).

(D) Quantitative analysis of HLA-G expression intensity in total CT29 EVTs obtained by exposure to laminin (n=2592) or laminin in the presence of a PKC$\alpha/\beta$ inhibitor, Gö6976 (n=2436). Analysis was performed in MATLAB and at least 2 biological replicates were used. (***p<0.0005). Data for laminin is same as used in Figure S3.

(E) Gene expression of *ITGA5, MMP2, ITGA1, MYC, CDH5, TP63, ELF5, HAND1, CDH1,* and *TEAD4* of EVTs obtained by exposure to laminin in the absence (blue) and presence (red) of the PKC$\alpha/\beta$ inhibitor, Gö6976 from CT29 hTSCs. Three biological replicates were used. (Error bars, S.E., *p<0.05 for comparison with undifferentiated hTSCs (dashed line), $^{\#}$p<0.05 for comparison with EVTs obtained by exposure with laminin at the same day).

(F) Confocal image of STB in the presence of a PKC$\alpha/\beta$ inhibitor, Gö6976 from CT29 hTSCs, staining for hCG and SDC-1. Nuclei were stained with DAPI.

(G) Confocal image of STB in the presence of a PKC$\alpha/\beta$ inhibitor, Gö6976 from SC102A-1 hTSCs, staining for hCG and SDC-1. Nuclei were stained with DAPI.

Scale bars are 100μm for all images.

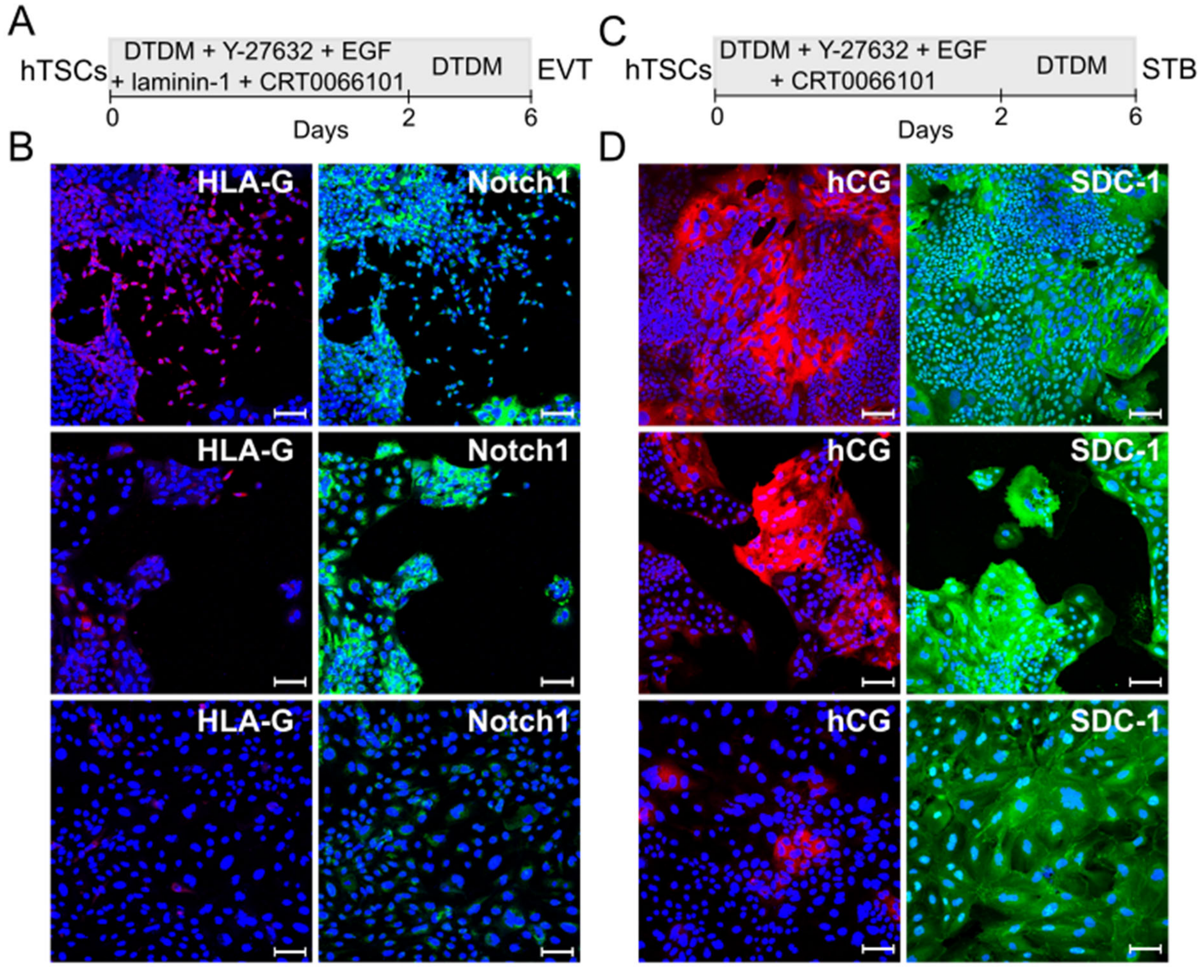


**Figure S5: Inhibition of PKD1 does not affect laminin-mediated EVT or STB differentiation.**

(A) Schematic of protocol for hTSC differentiation to EVT obtained by exposure to laminin in the presence of a PKD1 inhibitor, CRT0066101.

(B) Confocal image of EVT obtained by exposure to laminin in the presence of a PKD1 inhibitor, CRT0066101 from CT30 (top), CT29 (middle) and SC102A-1 (bottom) hTSCs, staining for HLA-G and Notch1. Nuclei were stained with DAPI.

(C) Schematic of protocol for hTSC differentiation to STB in the presence of a PKD1 inhibitor, CRT0066101.

(B) Confocal image of STB in the presence of a PKD1 inhibitor, CRT0066101 from CT30 (top), CT29 (middle) and SC102A-1 (bottom) hTSCs, staining for hCG and SDC-1. Nuclei were stained with DAPI.

Scale bars are 100µm for all images.

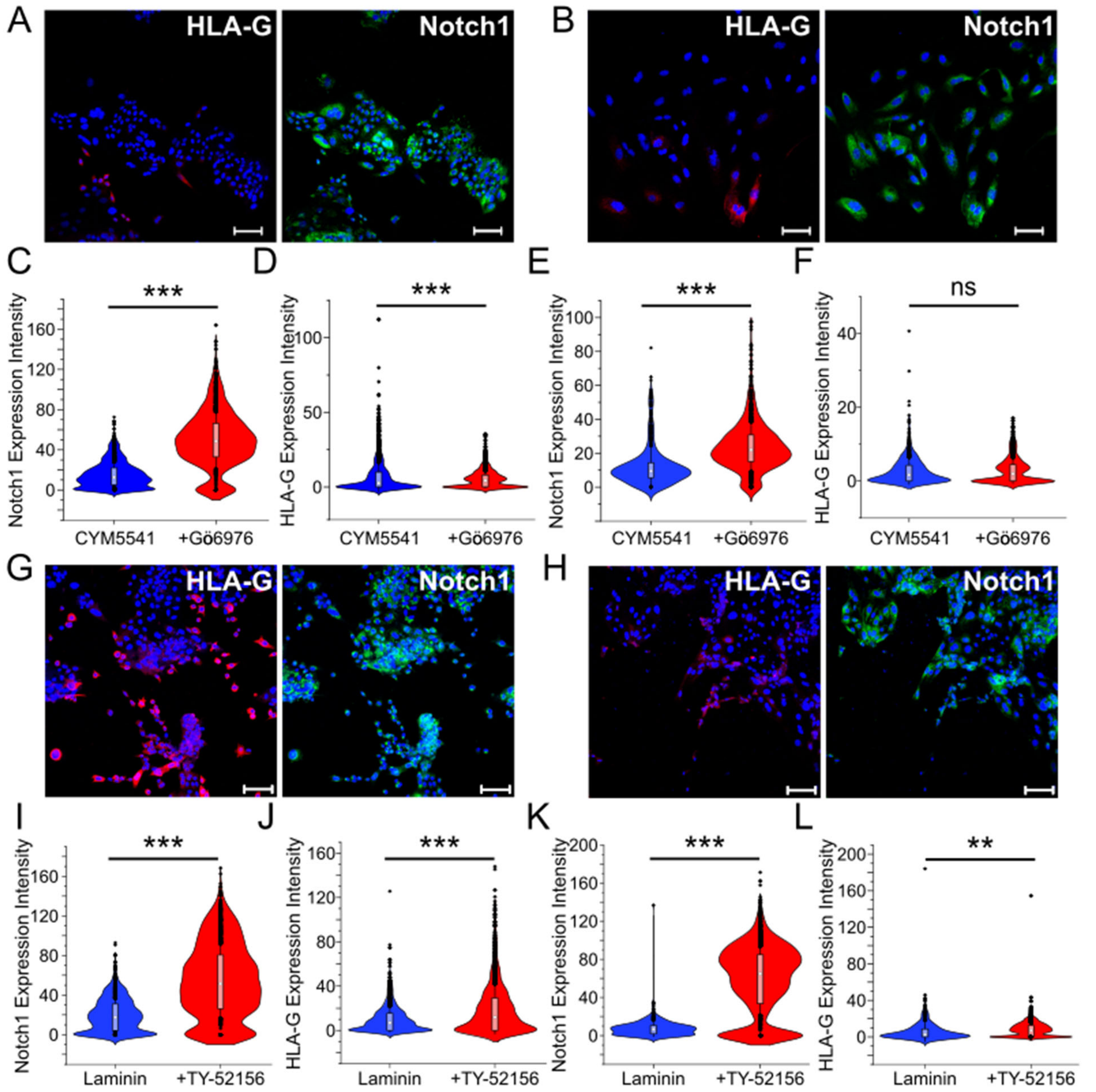


**Figure S6: PKCα/β inhibition differentially affects EVT differentiation mediated by laminin exposure and S1PR3 activation.**

(A) Confocal image of EVT obtained by exposure to a S1PR3 agonist, CYM5541 in the presence of a PKCα/β inhibitor, Gö6976 from CT29 hTSCs, staining for HLA-G and Notch1. Nuclei were stained with DAPI.

(B) Confocal image of EVT obtained by exposure to a S1PR3 agonist, CYM5541 in the presence of a PKCα/β inhibitor, Gö6976 from SC102A-1 hTSCs, staining for HLA-G and Notch1. Nuclei were stained with DAPI.

(C) Quantitative analysis of Notch1 expression intensity in CT29 EVTs obtained by exposure to a S1PR3 agonist, CYM5541 (n=4398) or CYM5541 in the presence of a PKC$\alpha/\beta$ inhibitor, Gö6976 (n=1984). Analysis was performed in MATLAB and at least 2 biological replicates were used. (***p<0.0005).

(D) Quantitative analysis of HLA-G expression intensity in CT29 EVTs obtained by exposure to a S1PR3 agonist, CYM5541 (n=4398) or CYM5541 in the presence of a PKC$\alpha/\beta$ inhibitor, Gö6976 (n=1984). Analysis was performed in MATLAB and at least 2 biological replicates were used. (***p<0.0005). Data for CYM5541 is same as used in Figure 3.

(E) Quantitative analysis of Notch1 expression intensity in SC102A-1 EVTs obtained by exposure to a S1PR3 agonist, CYM5541 (n=749) or CYM5541 in the presence of a PKC$\alpha/\beta$ inhibitor, Gö6976 (n=1340). Analysis was performed in MATLAB and at least 2 biological replicates were used. (***p<0.0005).

(F) Quantitative analysis of HLA-G expression intensity in SC102A-1 EVTs obtained by exposure to a S1PR3 agonist, CYM5541 (n=749) or CYM5541 in the presence of a PKC$\alpha/\beta$ inhibitor, Gö6976 (n=1340). Analysis was performed in MATLAB and at least 2 biological replicates were used. (ns, not significant). Data for CYM5541 is same as used in Figure 3.

(G) Confocal image of EVT obtained by exposure to a laminin in the presence of a S1PR3 antagonist, TY-51256 from CT29 hTSCs, staining for HLA-G and Notch1. Nuclei were stained with DAPI.

(H) Confocal image of EVT obtained by exposure to laminin in the presence of a S1PR3 antagonist, TY-51256 from SC102A-1 hTSCs, staining for HLA-G and Notch1. Nuclei were stained with DAPI.

(I) Quantitative analysis of Notch1 expression intensity in CT29 EVTs obtained by exposure to laminin (n=2592) or laminin in the presence of a S1PR3 antagonist (n=2040). Analysis was performed in MATLAB and at least 2 biological replicates were used. (***p<0.0005). Data for laminin is same as used in Figure S4.

(J) Quantitative analysis of HLA-G expression intensity in CT29 EVTs obtained by exposure to laminin (n=2592) or laminin in the presence of a S1PR3 antagonist, TY-51256 (n=2040). Analysis was performed in MATLAB and at least 2 biological replicates were used. (***p<0.0005). Data for laminin is same as used in Figures S3 and S4.

(K) Quantitative analysis of Notch1 expression intensity in SC102A-1 EVTs obtained by exposure to laminin (n=498) or laminin in the presence of a S1PR3 antagonist (n=5004). Analysis was performed in MATLAB and at least 2 biological replicates were used. (***p<0.0005). Data for laminin is same as used in Figure S4.

(L) Quantitative analysis of HLA-G expression intensity in SC102A-1 EVTs obtained by exposure to laminin (n=498) or laminin in the presence of a S1PR3 antagonist, TY-51256 (n=5004). Analysis was performed in MATLAB and at least 2 biological replicates were used. (**p<0.005). Data for laminin is same as used in Figure S3 and S4.

Scale bars are 100μm for all images.
